# Rotary substates of mitochondrial ATP synthase reveal the basis of flexible F_1_-F_o_ coupling

**DOI:** 10.1101/543108

**Authors:** B. J. Murphy, N. Klusch, J. Langer, D. J. Mills, Ö. Yildiz, W. Kühlbrandt

**Affiliations:** Department of Structural Biology, Max Planck Institute of Biophysics, Max-von-Laue Str. 3, Frankfurt 60438, Germany; Department of Molecular Membrane Biology, Max Planck Institute of Biophysics, Max-von-Laue Str. 3, Frankfurt 60438, Germany

## Abstract

F_1_F_o_-ATP synthases play a central role in cellular metabolism, making the energy of the proton-motive force across a membrane available for a large number of energy-consuming processes. We determined the single-particle cryo-EM structure of active dimeric ATP synthase from mitochondria of *Polytomella sp*. at 2.7-2.8 Å resolution. Separation of 13 well-defined rotary substates by 3D classification provides a detailed picture of the molecular motions that accompany *c*-ring rotation and result in ATP synthesis. Crucially, the F_1_ head rotates along with the central stalk and *c*-ring rotor for the first ∼30° of each 120° primary rotary step. The joint movement facilitates flexible coupling of the stoichiometrically mismatched F_1_ and F_o_ subcomplexes. Flexibility is mediated primarily by the interdomain hinge of the conserved *OSCP* subunit, a well-established target of physiologically important inhibitors. Our maps provide atomic detail of the *c*-ring/*a*-subunit interface in the membrane, where protonation and deprotonation of *c*-ring *c*Glu111 drives rotary catalysis. An essential histidine residue in the lumenal proton access channel binds a strong non-peptide density assigned to a metal ion that may facilitate *c*-ring protonation, as its coordination geometry changes with *c*-ring rotation. We resolve ordered water molecules in the proton access and release channels and at the gating aArg239 that is critical in all rotary ATPases. We identify the previously unknown *ASA10* subunit and present complete *de novo* atomic models of subunits *ASA1-10*, which make up the two interlinked peripheral stalks that stabilize the *Polytomella* ATP synthase dimer.

**One Sentence Summary:** Mechanisms of proton translocation and flexible F_1_-F_o_ coupling by rotary substates in a mitochondrial ATP synthase dimer

## Main Text

Mitochondria carry out controlled oxidation of reduced substrates to generate an electrochemical gradient across the inner mitochondrial membrane. Movement of protons down the gradient through the F_1_F_o_-ATP synthase drives the production of the soluble energy carrier ATP by rotary catalysis. The ATP synthases consist of two connected nanomotors: the membrane-embedded F_o_ subcomplex where proton translocation generates torque, and the soluble F_1_ subcomplex where the torque powers ADP phosphorylation (for a recent review, see (*1*)). In F_o_, a ring composed of 8-17 c-subunits, each containing a carboxylate ion-binding site, rotates against a bearing formed by a near-horizontal helix hairpin of the stationary *a*-subunit (*2*). Two aqueous half-channels at the interface of the *c*-ring and *a*-subunit allow protons to enter and leave the protonation site; upon protonation from the lumenal channel, a *c*-ring subunit travels by an almost full rotation through the hydrophobic membrane environment before releasing the proton in the matrix channel. In the membrane, the channels are separated by a mere 6 Å (*3*), with a strictly conserved arginine residue facilitating the passing of deprotonated, but not protonated, *c*-ring subunits. The energy from down-gradient proton translocation powers rotation of the *c*-ring and the firmly attached central stalk (subunits *γ,δ,ε*), which conveys rotation generated in F_o_ to power ATP synthesis. A two-helix bundle of the *γ*-subunit extends deep into the *α_3_β_3_* hexamer of the F_1_ head. Rotation of the central stalk induces conformational changes of subunits *α* and *β* that result in ADP phosphorylation. The two-domain oligomycin-sensitivity conferring protein, *OSCP*, connects the F_1_ head to a membrane-anchored peripheral stalk stator, preventing unproductive rotation of F_1_.

The dimeric mitochondrial ATP synthase from the chlorophyll-less unicellular alga *Polytomella* sp. contains the signature ATP synthase subunits *α_3_, β_3_, γ, δ, ε, c_10_, a*, and *OSCP*, while peripheral stalk and other membrane protein subunits typical of mammalian and fungal ATP synthases (*1*) are replaced in *Polytomella* and related species by proteins known as ATP-synthase associated (ASA) proteins, which bear no homology to other known ATP synthase components (*4, 5*). These subunits form a bulky peripheral stalk that links the two ATP synthase complexes into a stable dimer.

ATP synthase must couple translocation of eight to seventeen protons (*1*), depending on *c*-ring size, with phosphorylation of three molecules of ADP in the catalytic *β*-subunit sites of the F_1_ head. The stoichiometric mismatch of F_1_ and F_o_ poses a challenge to efficient energy conversion in the complex (*6, 7*). It has been proposed that the central stalk mediates flexible coupling between F_1_ and F_o_ (*8, 9*). Some authors suggest energy is stored by coiling of the *γ*-subunit two-helix bundle (*10*); others propose that energy is stored at the ‘foot’, where *γ* and *ε* are attached to the *c*-ring (*9*). Recent cryo-EM studies resolving rotary states of mitochondrial (*11*), bacterial (*12*) and chloroplast (*13*) ATP synthase have all found that the central stalk rotates as a rigid body. Substantial flexibility in the peripheral stalk is observed between rotary states (*11–14*), which suggests that peripheral stalk bending contributes to elastic energy storage. The robust, rigid peripheral stalks of the *Polytomella* ATP synthase make it an ideal system for examining the dynamics revealed by rotary states and substates by high-resolution cryo-EM, and for testing the idea that peripheral stalk flexibility is required for ATP synthase function.

## High-resolution map of a complete F-type ATP synthase dimer

Cryo-EM single-particle analysis of purified and active (Figure S1) *Polytomella* ATP synthase dimer yielded a map at 2.94 Å overall resolution (Table S1). Local masking improved the resolution to 2.7-2.8 Å in overlapping regions covering the full complex (Figure S1, S2, S3; Table S2, Movie S1). Symmetry expansion allowed each monomer to be treated independently in image analysis, and 3D classification of the central stalk and F_1_ head enabled separation of three primary rotary states, at resolutions of 2.8-2.9 Å in the F_1_ head and rotor (Figure 1). The nucleotide-binding sites *β_TP_* and *β_DP_* contain Mg^2+^ADP (Figure 1b,c,d) and *β_E_* is empty; this is the ADP-bound complex, which is to be expected as it was prepared under substrate-depleted conditions. The ‘arginine finger’ (*15*) of aArg429 extends toward the nucleotide phosphate in *β*_DP_ but away from it in *β*_TP_ (Figure 1c,d). Our map provides the highest available resolution of a complete ATP synthase. Cryo-EM data were complemented by genomic sequencing and mass spectrometry analysis of the purified protein, enabling us to build a full atomic model of the 1.58 MDa complex, with 62 copies of 18 different subunits and a total of 14,240 residues fitted. (Figure 1; Figure S4; Table S3). The model includes the completed polypeptide sequence of subunits *ASA2* and *ε* and the previously unknown subunit *ASA10*.

**Figure 1.**
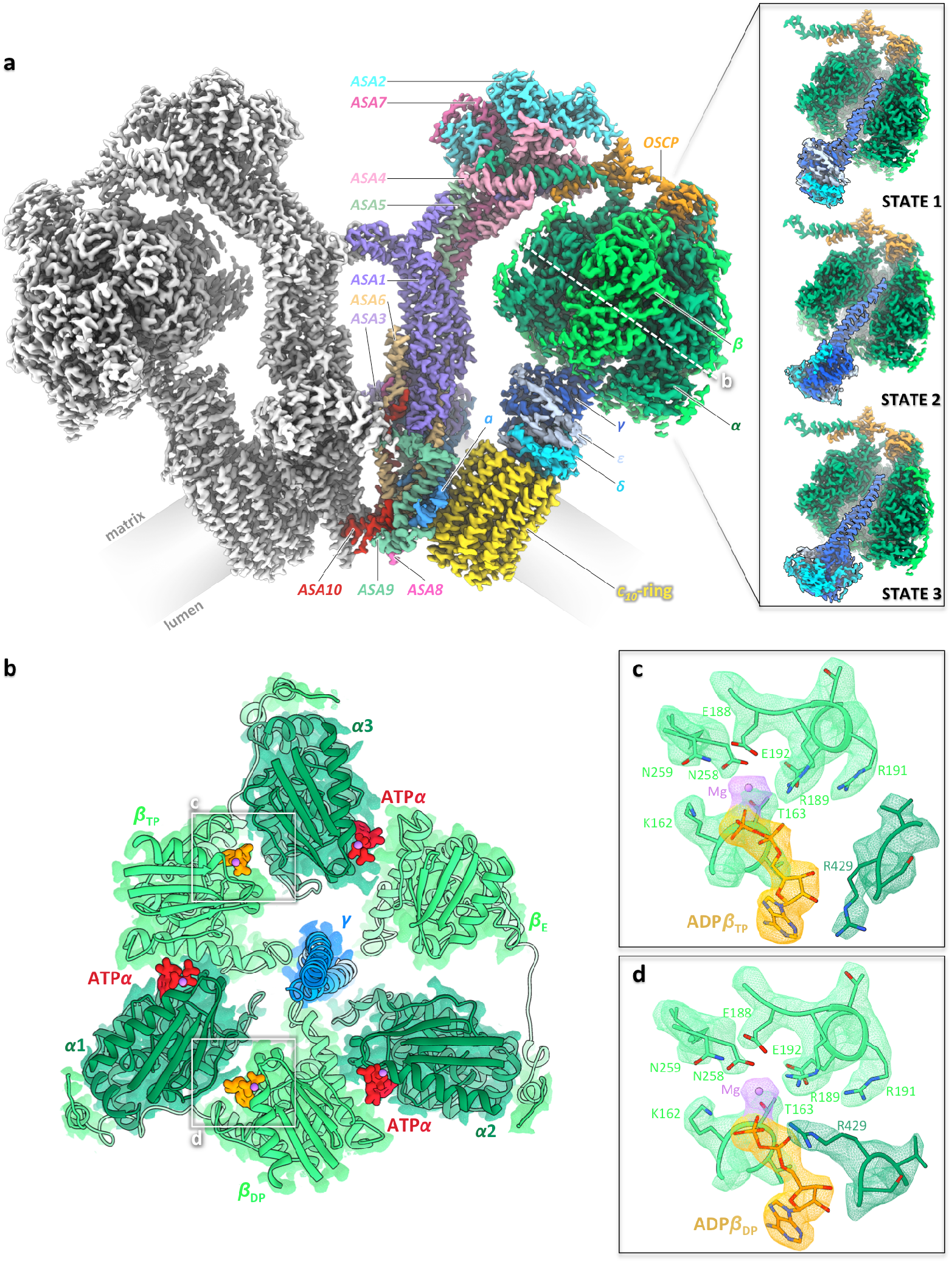
High-resolution structure of the mitochondrial F_1_F_o_ ATP synthase dimer from *Polytomella* sp. **a)** Composite cryo-EM map density of the 62-subunit, 1.58 MDa *Polytomella* dimer, colored by subunit on the right. The *c_10_* rotor ring (yellow) and subunit *a* (bright blue) make up the F_o_ motor complex. Proton translocation through F_o_ causes rotation of the *c*-ring and the attached central stalk subunits *γ*(dark blue), *δ* (cyan) and *e* (light blue). The F_1_ head consists of three catalytic *β*subunits (bright green) and three *a* subunits (dark green). Long C-terminal extensions of *β*wrap around the *a* subunits on the outside of F_1_ (see Movie S5). The two-domain *OSCP* subunit (orange) links the three *a* subunits to the peripheral stalk subunits *ASA1-10* (see Figure 4). Subunit *ASA10* (red) connects the two F_1_F_o_ monomers in the membrane (see Figure S7, S9). The local map resolution is 2.7 Å for the peripheral stalk, F_o_ and *c*-ring, and 2.8 to 2.9 Å for the F_1_ head and central stalk (see Figure S2). **Inset:** Three primary rotary states 1, 2 and 3 of the F_1_F_o_ monomer are related by ∼120° rotation of the central stalk within the *α_3_β_3_* assembly. Unless otherwise specified, subunit coloring is consistent throughout all figures. **b)** Section through F_1_ at the level of bound nucleotides. *β*_DP_ and *β* sites contain ADP (orange) and *β*_E_ is empty. The three a-subunits bind a structural ATP (red). Nucleotide-coordinating Mg^2+^ ions are violet. The central stalk subunit *γ* engages with the catch loop of the *β*_E_ subunit (see Figure S5). **c,d)** Catalytic sites of subunits *β*_DP_ and *β*_TP_. A well-defined arginine sidechain of subunit *α*(the “arginine finger”) extends toward the nucleotide phosphate in the *β*_DP_ site. ATP in the *β*_TP_ site has hydrolyzed to ADP during protein isolation.

## Rotary substates reveal concerted rotation of F_1_ and central stalk

To describe the broad range of thermally-accessible motion of F_1_F_o_-ATP synthases, we sorted the symmetry-expanded dataset into 12 classes. Further subdivision of a mixed class yielded a total of 13 unique rotary substates (Figure S2) which were refined to 2.8-4.2Å resolution (Figure S3; Table S2). The subclasses can be placed into a meaningful sequence according to the rotational position of the rotor with respect to the stator. Each subclass was assigned to one of three primary rotary states (Figure S1), with six subclasses for state 1 (1A-1F) (Figure 2), four for state 2 (2A-2D) and three for state 3 (3A-3C). The 3 primary states are separated from one another by ∼120° rotation of the central stalk within the F_1_ head (Figure 1a, inset; Figure S2); in contrast, substates of any given rotary state differ by concerted rotation of the F_1_ head and rotor within the first 15° to 32° of each ∼120° step (Figure 2; Figure 3; Movie S2, S3). Contact between the *β_E_* site and the *γ*-subunit (the ‘*β*-catch loop’ (*16*)) is maintained between substates (Figure S5; Movie S2), as is the nucleotide occupancy of *β*-subunits (Figure 3b). The pivot point of this movement is the single peptide chain connecting the two *OSCP* domains (henceforth the *‘OSCP* hinge’) (Figure 4; Movie S4), while the top of the peripheral stalk flexes only slightly. In comparing all 13 substates, no coiling of the central stalk is apparent, but the extreme C-terminus of *γ* bends slightly in some substates.

**Figure 2.**
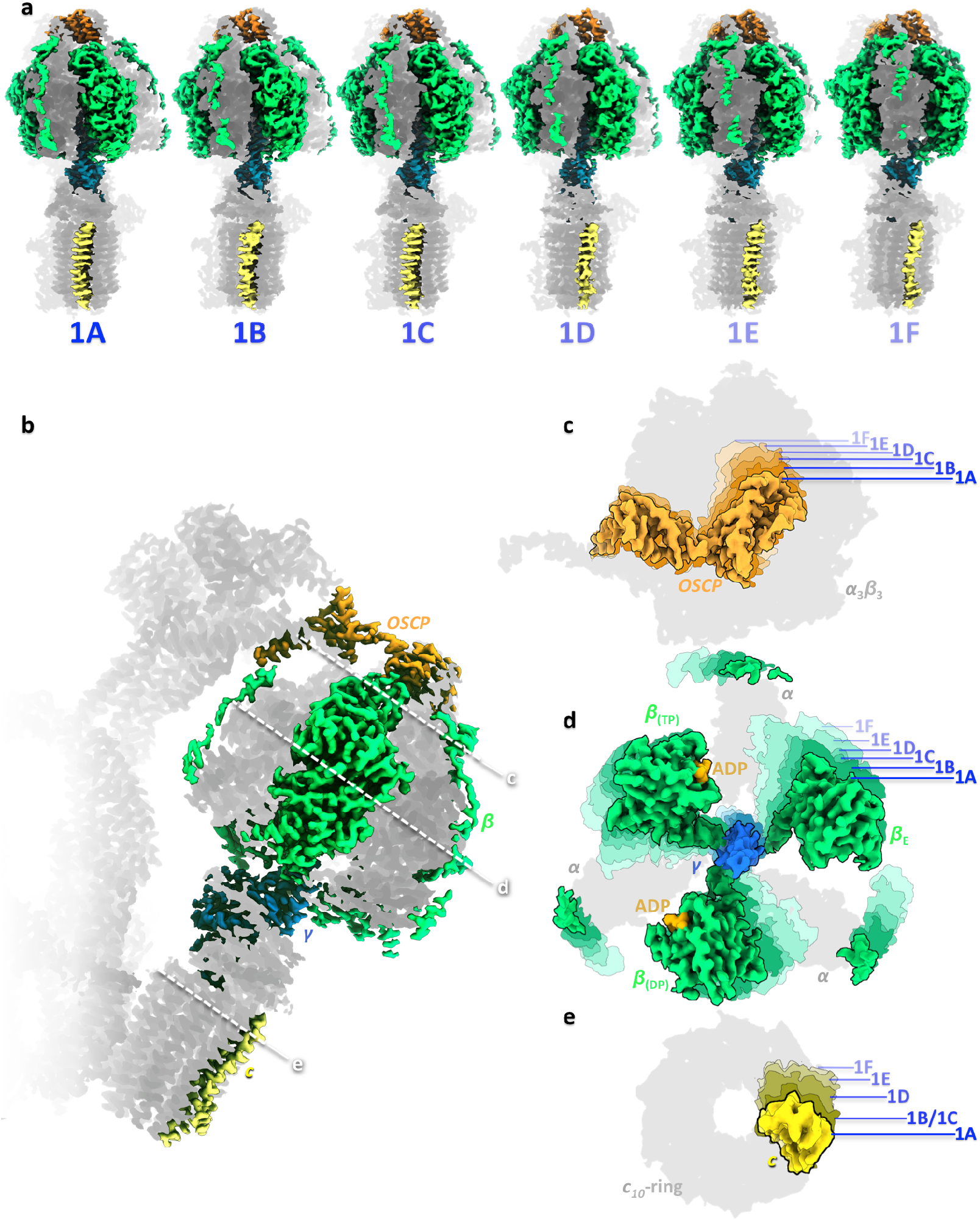
Rotary substates of mitochondrial ATP synthase. The three primary rotary states are subdivided into a total of 13 rotary substates at resolutions between 2.8 and 4.2 Å (see Figure S1 and Table S2). The number of resolved substates and angular increments between them differ for each of the three primary rotary states. **a)** Density maps of rotary substates 1A to 1F ordered in ATP synthesis direction according to the position of the central stalk with respect to the peripheral stalk. **b)** Composite cryo-EM density map of rotary substate 1A at 2.9 Å resolution. **c-e)** An overlay of substate maps indicates a concerted movement of *OSCP* (orange, **c)**, the F_1_ head (green, **d)**, *γ*(blue, **d)**, and the *c*-ring (yellow, **e)** from substate 1A to 1F. Projected map densities of other subunits are shown in light grey for reference.

**Figure 3.**
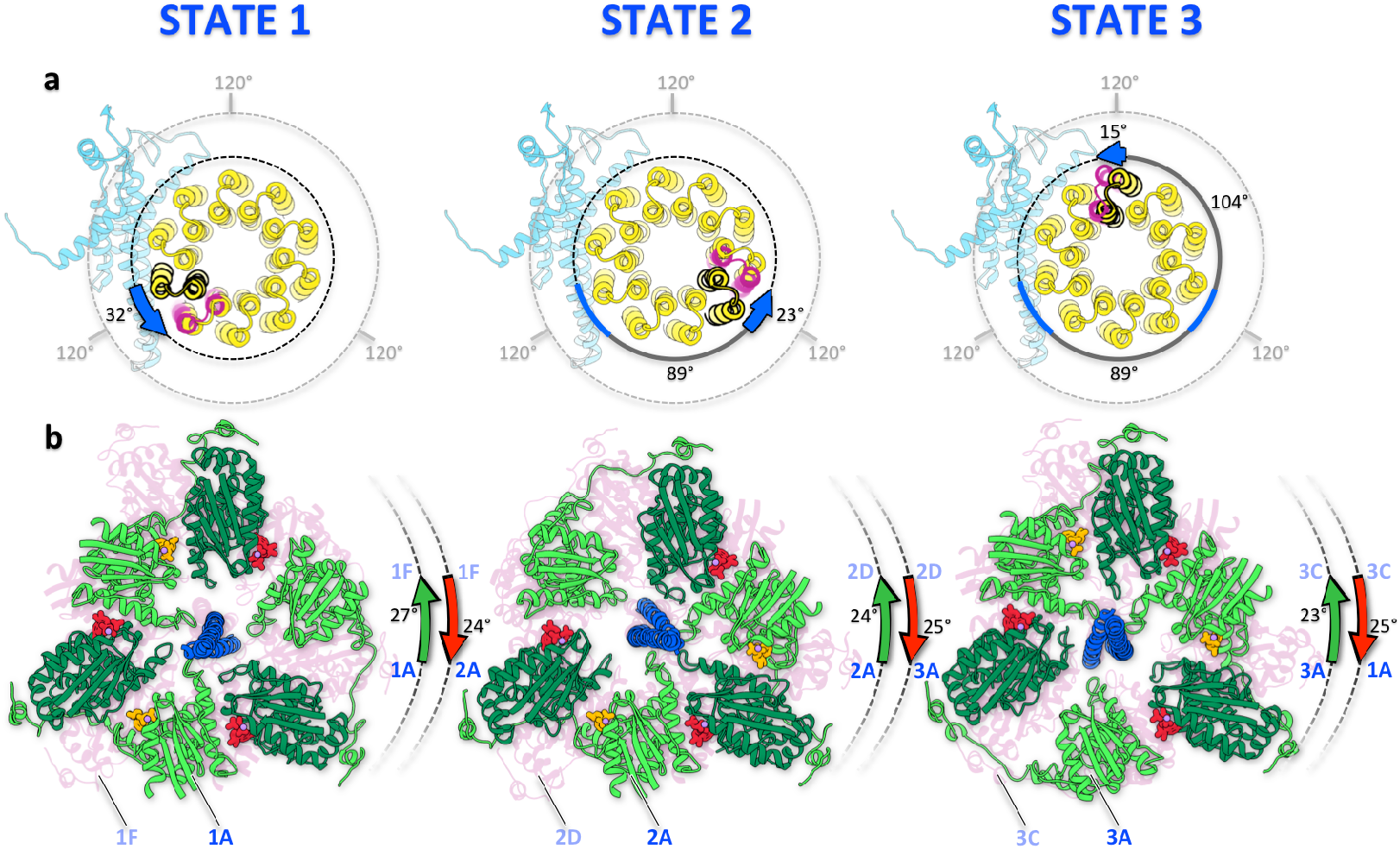
Concerted rotation of F_1_ with central stalk and *c*-ring. **a)** Progressing in ATP synthesis direction, the *c_10_*-ring rotates counter-clockwise (as seen from F_1_) with respect to the σ-subunit, by up to 32° for substates of the same primary rotary state. The primary rotary states differ by power strokes of ∼120°. **b)** Between substates of a given rotary state, the F_1_ head rotates together with the *c*-ring and central stalk before recoiling to its original position in the first substate of the subsequent primary rotary state.

It is clear that in the first ∼30° of rotary steps 1 and 2, the *c*-ring advances by almost one subunit with respect to the a-subunit in moving between substates (Figure 3a), while the position of *γ* with respect to the *α_3_β_3_* hexamer remains virtually unchanged (Figure 3b; Figure S5). Between substates of state 3, the *c*-ring advances by about half a subunit (Figure 3a). As the rotor moves a further ∼90° to the starting position of the next primary rotary state (e.g. from state 1F to 2A), the F_1_ head recoils by ∼30° to its original position, for a cumulative ∼120° power stroke of *γ* within F_1_ for this step. Given that this motion is thermally accessible for the complex at equilibrium, it would almost certainly contribute to flexible coupling of F_1_ and F_o_ also under turnover conditions.

The interaction between F_1_ and *γ* appears strongest in the catch loop region (*16*) of *β_E_*, where conserved residues *β* Asp318, *β*Asp321, and *β*Thr320 (*β*Asp302, *β*Asp305, *β*Thr304 in *E. coli*) form ionic and hydrogen-bond interactions with *γ*Arg296 and *γ*Gln297 (*Ec γ*Arg268 and *γ*Gln269). The salt bridge between *β*Asp321 and *γ*Arg296 forms via a resolved water molecule (Figure S5). Throughout the 20°-30° concerted rotation of F_1_ with *γ*, the interaction between the *β_E_* catch loop and *γ* is maintained, although subtle loop movements suggest that the hydrogen-bonding partner of *β*Asp318 changes from *γ*Gln297 to *γ*Asn293 between substates (*e.g*. from state 1A to 1F; Figure S5). Disruption of these interactions by mutation in *E. coli* eliminated or strongly reduced ATP hydrolysis activity and the ability to grow on succinate (*16*). A proposed function of such an interaction is that, in ATP synthesis mode, it would stall rotation of *γ* until the *β_E_* site has bound ADP, to prevent unproductive rotation (*16*), although the dramatic impact of catch loop mutations on both synthesis and hydrolysis activity implies that this region plays a broader role in the conformational changes required for catalysis. Our findings make it clear that this interaction does not stall rotation completely; rather, a proton may be translocated at F_o_ while the F_1_ head waits for substrate binding.

Studies aiming to characterize flexible coupling have focused on the central stalk (*17*), often working with an F_1_ and rotor subcomplex. When the whole F_1_F_o_ complex was examined, it was nonetheless normally attached to substrate and reporter molecules at subunits *β* and *c* to probe for flexibility along the F_1_-F_o_ axis (*9,18,19*). A recent single-molecule study of F_1_F_o_ reported forward and backward stepping of the rotor by ∼30°, i.e. up to one c-subunit (*18*). However, with F_1_ anchored to the support by subunit *β* and a probe attached to *c*, flexibility at *OSCP* would not be detected. The observed ∼30° stepping angle appears to reflect regular energy minima in the *c*-ring/*a*-subunit interaction rather than joint rotation of the *c*-ring and central stalk with F_1_. Single-molecule studies of flexible coupling may need to explore a wider range of anchoring points.

Our structure indicates that bending at the *OSCP* hinge (Figure 4; Movie S4), by mediating flexible coupling between F_1_ and F_o_, plays an important mechanistic role in ATP synthesis. A body of literature suggests that this process has broad physiological implications, with a number of regulatory mechanisms acting on *OSCP*. The regulator of mitochondrial permeability transition, cyclophilin D, binds to *OSCP* and inhibits ATP synthase activity (*20*), as does the immunomodulator Bz-423, which affects both V_max_ and K_M_ for ATP hydrolysis (*21*). Acetylation of a conserved lysine near the hinge region of *OSCP* in response to exercise stress affects mitochondrial ATP levels (*22*). The hormone estrogen binds to *OSCP* (*23*) and inhibits ATP synthase activity (*24, 25*) with possible relevance to neuroprotection (*26*). The tumor suppressor p53, under non-stress conditions, binds to *OSCP* and promotes increased O_2_ consumption and decreased ROS levels in mitochondria (*27*). These diverse and potent modulations of ATP synthase activity cannot be reconciled with a model of ATP synthase in which *OSCP* forms a static bridge between the F_1_ head and peripheral stalk. Our results show that, on the contrary, *OSCP* plays a dynamic role in the flexible coupling of F_o_ and F_1_. The concept of a dynamic *OSCP* will be essential to understand and exploit modulation of ATP synthase activity, with relevance to neurodegeneration and traumatic brain injury, ischemia-reperfusion injury, and immunomodulation, among others.

**Figure 4.**
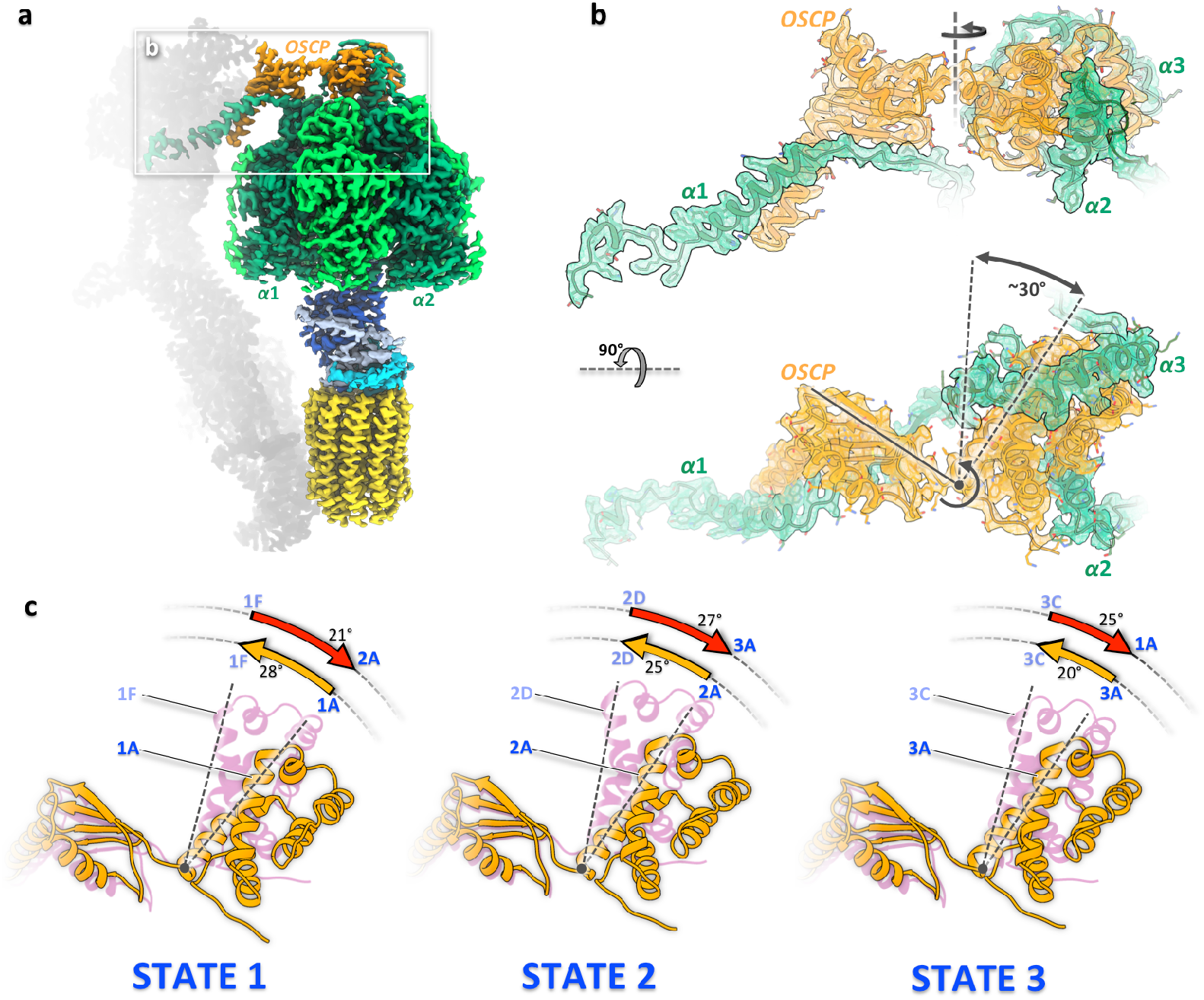
Subunit *OSCP* connects the F_1_ head and peripheral stalk as a flexible hinge. **a)** Overview of F_1_F_o_ monomer indicating the position of subunit *OSCP* (orange) on top of the F_1_ head (green) relative to the peripheral stalk (gray). **b)** The two-domain *OSCP* interacts with the helical C-terminal extensions of subunits *α_1_, α_2_* and *α_3_* (green) (see Movie S4). Extensions of *α_2_* and *α_3_* form short helix bundles with helices of the globular N-terminal *OSCP* domain, while *α_1_* binds to the elongated C-terminal *OSCP* domain that is attached to the peripheral stalk via the *α*_1_ extension. The two *OSCP* domains are connected by a flexible peptide link that facilitates ∼30° back and forth rotation of the N-terminal domain with F_1_ (black arrows) relative to the C-terminal *OSCP* domain and peripheral stalk. **c)** Between substates of a given primary rotary state, the N-terminal *OSCP* domain rotates with F_1_ by up to 28° (straight dashed lines) in synthesis direction (orange arrows) and then recoils to its starting position in the first substate of the next primary rotary state (red arrows).

Several lines of evidence support the idea that the mechanism of flexible coupling is conserved across all types of rotary ATP synthases: 1) the structure and polypeptide sequences of *β, γ*, and *OSCP* subunits involved in this process are highly conserved; 2) a canonical *OSCP* (or *δ* in chloroplasts and bacteria) is found in all three lineages of ATP synthase known to have independently acquired novel peripheral stalk components (*28, 29*); 3) a cryo-EM study of bovine ATP synthase (*11*) identified rotary substates of this complex, albeit at lower resolution, indicating rotation of F_1_ relative to the remainder of the complex. Re-evaluating these results in light of our data, they do in fact show concerted rotation of F_1_ with the central stalk; 4) well-known inhibitors and regulators of ATP synthase bind to *OSCP*, affecting the catalytic rate and K_M_ (*21, 30*), suggesting a mechanistic rather than merely structural role for this subunit.

## The essential His-248 ligates a metal ion that is sensitive to *c*-ring rotation

In respiring mitochondria, the pH of the lumen is 7.2 (*31*), while the pK_a_ of the *c*-ring carboxylate in an aqueous environment is around 4.5 (*32*); thus, protonation of the *c*-ring needs to be assisted by the local environment in the lumenal entrance channel in order to achieve high turnover rates. Mutational studies in *E. coli* have established that the well-conserved lumenal channel environment is necessary for activity, with residues *a*Glu219 and *a*His245 essential for both ATP synthesis and proton permeability of F_o_ in membranes stripped of F_1_. The mutations *a*E219H and *a*H245E show low levels of activity, while the double mutant is more active than either single mutant (*33*), suggesting that the residues interact directly to facilitate *c*-ring protonation. Our map shows a strong nonpeptide density ligated by *a*His248 (*a*Glu219 in *E.coli*) and *a*His252 of *a*H5 (Figure 5) with bond lengths of 2.1 - 2.2Å. The strong density and coordination environment favor assignment to a metal ion, provisionally annotated as Zn^2+^, on account of its abundance, flexible coordination geometry and lack of redox activity. At high threshold levels the bound species appears diatomic, perhaps due to a heavy ligand (Figure 5b, f-h). Both ligating His residues are present in mammalian ATP synthases (Figure S6). The aHis248 position is occupied by His or Glu in all F-type ATP synthases, with the notable exception of Na^+^-translocating complexes (Figure S6). Given its likely role in protonating the *c*-ring, this site thus appears to contribute to selectivity for H^+^ vs. Na^+^, in addition to known differences in the *c*-ring binding site (*34*). Recently published high-resolution cryo-EM maps of yeast mitochondrial (*35*) and spinach chloroplast ATP synthase (*13*) both indicate a strong, unattributed non-peptide density at the same position in the lumenal channel, near the residue equivalent to *a*His248 (*a*His185 and *a*Glu198, respectively) (Figure S6). In both instances, a single strong density was observed as compared to the two densities in our map, which may reflect the lower local resolution (3.6 Å and 3.4 Å) of the yeast and spinach maps.

**Figure 5.**
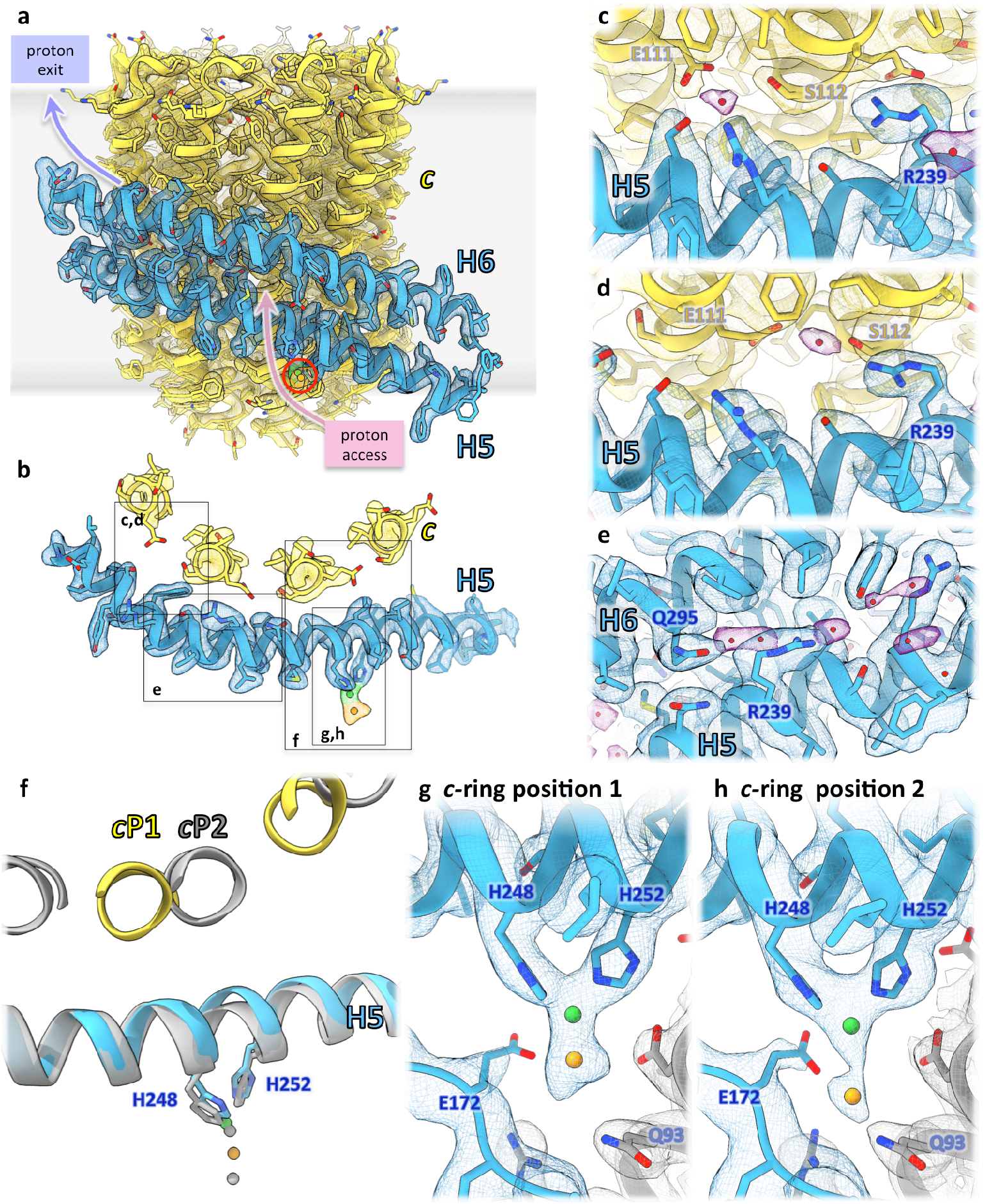
Metal ion and ordered water molecules in the F_o_ proton access and release channels. **a)** The H5/H6 helix hairpin of subunit *a* (light blue) forms the bearing against which the *c_10_*-ring (yellow) rotates, shown here for *c*-ring position 1. **b)** Sectional view of H5 and outer-ring c-subunit helices with proton-binding cGlu111, *c*-ring position 1. **c,d)** Water molecules (purple density) coordinated by the ion-binding cGlu111 and adjacent cSer112 in the matrix channel for *c*-ring positions 1 (**c**) and 2 (**d**). Maps are displayed at density thresholds of 0.035 for subunit *a*, 0.025 for the *c*-ring, and 0.025 (**c**) or 0.018 (**d**) for water. **e)** The strictly conserved *a*Arg239 and *a*Gln295 that separate the proton access and release channels in the membrane bind two water molecules, shown in the concensus C2-refined map of the F_o_ region. **f-h)** Conserved residues *a*His248 and *a*His252 in H5 of subunit *a* coordinate a metal ion. Bond distances change depending on *c*-ring position. The predominant *c*-ring position (position 1, colored, and panel **h)** accounts for 58% of particles, while a second position, differing by rotation through roughly 13°, (position 2, gray, and panel **g)** accounts for 33%. The overlay of **g** and **h** in panel **f** indicates a small lateral displacement of H5 with its metal-coordinating sidechains. Except where specified, all maps in a given panel are rendered at the same density threshold.

Masked 3D classification of the *c*-ring and a-subunit yields two distinct *c*-ring rotary positions at 2.7Å and 3.1 Å resolution, representing 58% and 33% of particles (Figure S2). The positions differ by rotation of the *c*-ring by approximately one third of a *c*-ring subunit (13°) (Figure 5f). Comparing these maps, it is clear that the configuration of the metal ion in the lumenal channel changes with rotation of the *c*-ring, with the metal-ligand distance measuring 2.3 and 3.7Å for *c*-ring positions 1 and 2 (Figure 5f-h). Sidechains of *a*Glu172 and ASA6Gln93 coordinate the ligand in state 2, with approximate distances of 2.8 and 2.2 Å. We observe a slight but significant change in curvature of hairpin helix H5 between the two *c*-ring positions (Figure 5f), which is likely due to mechanical pressure of the *c*-ring subunit against the helix, with flexibility of H5 facilitated by the conserved *a*Gly247 residue. The simplest explanation of our observations is therefore that the rotating *c*-ring pushes on aH5, inducing a slight change in curvature, which in turn changes the coordination of the bound metal ion. In both positions, distances between the His*ε*N and metal ion are in the 2.1-2.2 Å range, consistent with dative bonds and thus with unprotonated His*ε*N. Available structural and mutational data strongly suggest that the role of *a*His248 is to protonate *a*Glu288, which would require that *a*His248 is itself transiently protonated. Protonation of either His or Glu at this position would preclude their ability to coordinate a metal ion. This suggests the existence of a third state in which *a*His248 is protonated, leaves the metal ion and approaches *a*Glu288 more closely than the distance *a*His248Nε – *a*Glu288O of 5.5 Å in our model, which is too far for direct proton transfer. If protonation of *a*Glu288 by *a*His248 is very fast, the structure of this state would not be resolved.

Although further work will be needed to clarify the identity, dynamics, and function of this metal ion, the following working model may help to direct future studies: physical pressure of the *c*-ring on aH5 stretches the aHis248-metal ion bond, favoring its protonation. Upon protonation aHis248 leaves the metal ion and transfers a proton to aGlu288, which is then passed on to the *c*-ring glutamate, most likely via a water, since the residues do not appear to be able to make direct contact at any rotary position of the *c*-ring. Such a mechanism would synchronize proton delivery with *c*-ring rotation.

## Ordered water in the aqueous half-channels and at the essential Arg-239

The membrane-embedded a-subunit and *c*-ring together define the pathway for protons to move across the inner mitochondrial membrane (Figure 5a). The sequences of both subunits are very similar to those of other ATP synthases, even those of bacteria and chloroplasts (*1*). Previously we showed (*3*) that the aqueous half-channels that conduct protons to and away from the *c*-ring glutamate are separated by a distance of 5-7 Å in the centre of the membrane. We now present a high-resolution structure of this critical region. Our map shows ordered water molecules within both channels (Figure 5c-e). In the matrix channel, a water molecule is coordinated by cSer112 opposite the ion-binding cGlu111 of the adjacent c-subunit, favoring the idea that the *c*-ring glutamate may be directly deprotonated by water in this location. An interesting finding is the density of two water molecules enclosed by the strictly conserved *a*Arg239 and *a*Gln295, in a pocket which appears not to be continuous with the aqueous channels (Figure 5e). The interaction of these highly conserved residues with each other via a water molecule would stiffen the *a*Arg239 sidechain. In the lumenal proton access channel the well-ordered water lies between the assigned metal ion density and the strictly conserved *a*Arg239. Examining these waters, it is clear that the channel extends to the *a/c* interface close to *a*Arg239, as previously reported (*3*). Although the *c*-ring glutamate is exposed to the aqueous environment immediately adjacent to *a*Arg239, the results of mutational studies of aHis245 and *a*Glu219 in *E. coli* suggest that direct protonation of the *c*-ring from water in the channel is rare; rather, the *c*-ring is likely protonated as it rotates past *a*Glu288 (*a*His245 in *E. coli*), immediately before passing into the hydrophobic lipid bilayer environment. It is therefore worthwhile to consider the role of water in the lumenal channel permeating all the way to *a*Arg239. Two explanations seem possible: 1) the aqueous environment primes the *c*-ring glutamate for protonation by inducing an outward-facing conformation (*32*) and/or 2) the small distance between half-channels, which would generate a substantial field and resulting torque on the *c*-ring glutamate, is needed to overcome the energetic barrier of moving the *c*-ring glutamate past the gating Arginine 239 (*1*)(*3*).

## Novel C-terminal β-extensions contact the adjacent α-subunit

In chlorophycean algae that include *Polytomella*, the mitochondrial catalytic *β*-subunit resembles those of the canonical ATP synthases closely, except that the chlorophycean subunit has acquired a ∼60-residue C-terminal extension. Our map shows that this extension wraps vertically around the adjacent α-subunit (Movie S5) and moves with subunit *β* as it adopts an open or closed conformation. A ∼15-residue N-terminal extension of the *α*-subunits unique to chlorophycean algae is in a different configuration for each subunit (Figure 4b, Movie S4). One of these extensions, together with the distal *OSCP* domain, anchors one a-subunit to the peripheral stalk, while the other two form short helix bundles with the proximal globular domain of *OSCP*, attaching it to F_1_. The N- and C-terminal extensions would enhance the stability of the chlorophycean mitochondrial ATP synthase, which may be an advantage under stress conditions, as when *Polytomella* converts to a dormant cyst in nutrient-depleted environments (*36*).

## ATP-synthase associated proteins 1-10 form a rigid peripheral stalk

In *Polytomella*, dimer formation is mediated by the peripheral stalk consisting of ten *ASA* subunits that have no homologues in other known ATP synthases outside this class of green algae. We have modelled all nine previously described *ASA* subunits unambiguously into the 2.7 Å map (Figure 1; Figure S4; Movie S1), as well as an unknown subunit that we term *ASA10* (Figure S7). Mass spectrometry of the purified complex and DNA sequencing of the ∼50Mb *Polytomella* genome allowed us to identify *ASA10* (Figure S7) and to obtain complete sequences for subunits *ε* and *ASA2* (Figure S8). All subunits were built into the cryo-EM map *de novo*, as there are no homologous structures in the database.

The stability of the *Polytomella* dimer is due to the rigid peripheral stalks and their close interaction. *ASA1* is at the core of the soluble stalk region, and forms a bridge between monomers with a helix-turn-helix motif 75 Å above the membrane surface (Figure S9). F_o_ has six trans-membrane helices and four long, membrane-intrinsic helices which form the two helix hairpins of the conserved a-subunit (*2*). Of the six trans-membrane helices, two belong to *ASA6* and one each to *ASA5, ASA8, ASA9*, and *ASA10*. The two *ASA10* subunits form the primary dimer interface on the lumenal side of the membrane, and their C-termini are intertwined (Figure S7, S9). Little or no other direct interaction is seen within the membrane, with lipids or detergent filling the space between monomers. *ASA3* forms the characteristic Armadillo repeat domain above the membrane surface on the matrix side, and *ASA2, ASA4* and *ASA7* sit atop the peripheral stalk.

## Outlook

We have determined the structures of *Polytomella* ATP synthase in 13 rotary substates, allowing us to visualize most if not all of its thermally-accessible conformations. Critically, we have shown that the F_1_ head and rotor move together through a 20°-30° rotation; thus, the *c*-ring rotates relative to the a-subunit while the position of the central stalk within the F_1_ head is preserved (Figure 6). This movement ensures flexible coupling of the two symmetry-mismatched nanomotors of ATP synthase, F_o_ and F_1_, for all possible *c*-ring stoichiometries. Flexible coupling appears to be mediated predominantly by *OSCP*, which is emerging as an important target of cellular and pharmacological control. Whereas most previous structural studies of ATP synthase have characterized inhibited complexes, this work has instead examined the product-bound form of the complex, not autoinhibited and free of any added inhibitors, which may account for the increased conformational flexibility we observe. Further studies will address the energetics of *OSCP* bending, and integrate these results into a full catalytic scheme of ATP synthesis. A metal ion bound at the conserved residue aHis248 within the lumenal channel is sensitive to the *c*-ring rotary state and is likely to play an important role in protonating the *c*-ring carboxylate. An important unanswered question is how the strictly conserved aArg239 facilitates movement of deprotonated, but not protonated, *c*-ring subunits between the aqueous half-channels. The high-resolution detail provided by our model will be important for answering this question.

**Figure 6.**
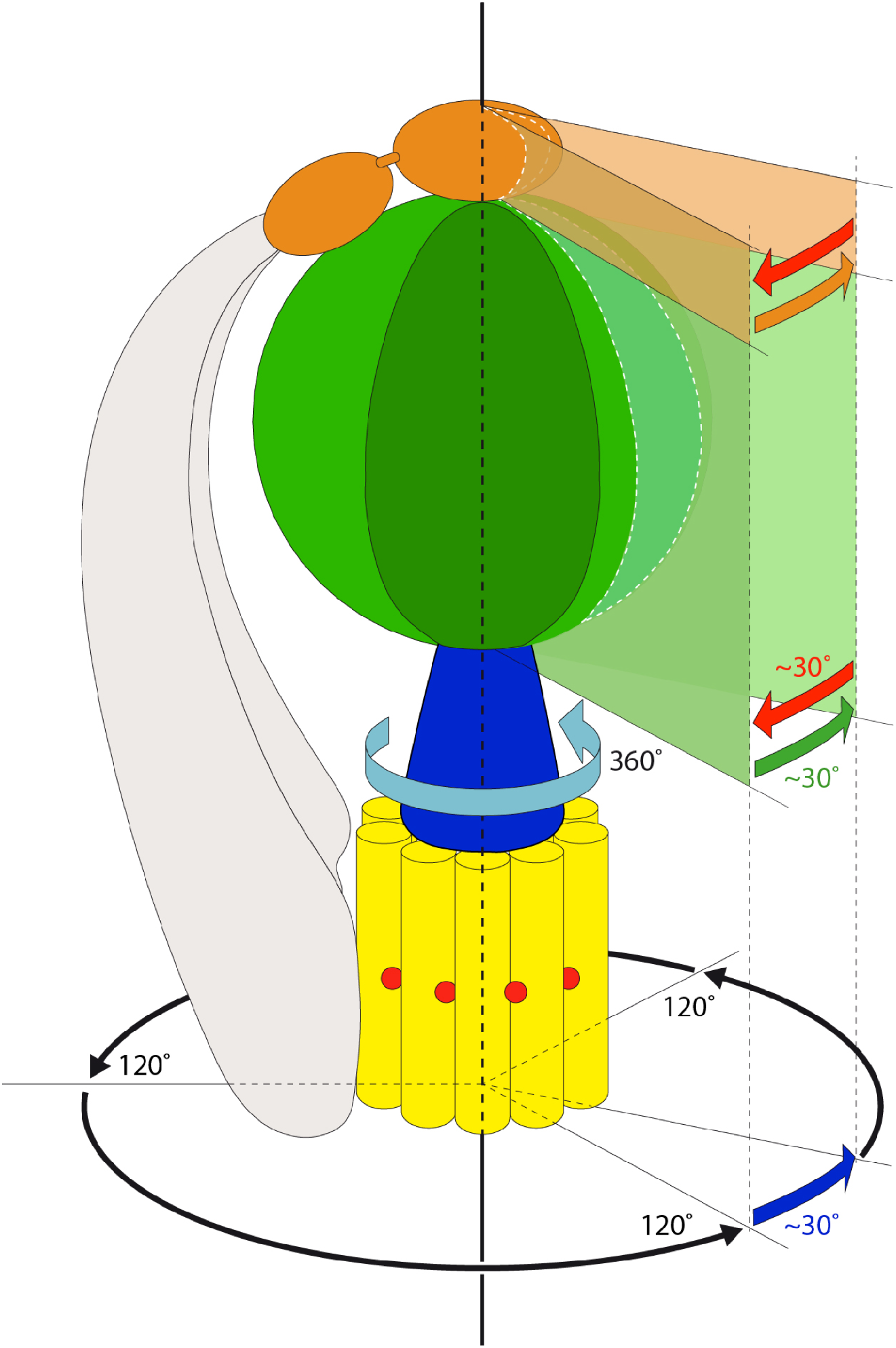
Concerted movement of F_1_ and rotor in F_1_F_o_ ATP synthase facilitates flexible coupling of F_o_ and F_1_. The F_1_ head (green) rotates together with the *c*-ring rotor (yellow) and central stalk (blue) for the first 20-30° of each ∼120° primary rotary step. *OSCP* (orange) forms a flexible link between the F_1_ head and the peripheral stalk (gray). F_1_ and the proximal *OSCP* domain then together recoil to their original position and the central stalk rotates ∼120° within F_1_ to the next primary rotary state. In this way, rotation of the central stalk within *α_3_β_3_* and rotation of the *c_10_*-ring in F_o_ are flexibly coupled.

## Acknowledgements

This work was funded by the Max Planck Society. BJM acknowledges funding from an EMBO Long-term fellowship (EMBO LTF 702-2016).

## Author contributions

WK initiated and supervised the study. NK grew *Polytomella* cultures, isolated, purified and analysed ATP synthase dimers biochemically. BJM and NK prepared cryoEM specimens and collected data. BJM performed image analysis and identified and analysed rotary substates. JL performed mass spectrometry. DJM maintained cryoEM instrumentation and EM alignment. BJM, NK and JL, ÖY, WK analysed data. BJM, NK and WK wrote the manuscript.

## Data and materials availability

Maps and models are publicly available through the EMDB and PDB databases (Tables S2 and S3). Mass spectrometry data are publicly available through the PRIDE database (accession number XXXX)(*37*).

## Supplemental Materials

Materials and Methods

Figures S1-S9

Tables S1-S3

Movies S1-S5

References 37-61

