## Supplemental methods, figures and tables for "Rotary substates of mitochondrial ATP synthase reveal the basis of flexible F_1_-F_o_ coupling"

### Supplementary Material

#### Materials and Methods

##### Protein isolation and purification

ATP synthase from *Polytomella sp.* was purified as described previously (1) with slight modifications. Mitochondrial membranes (175mg) were solubilized for 30min at 4°C in a volume of 12 ml buffer containing 30 mM Tris-HCl, pH 7.8, 2 mM $MgCl_2$, 50 mM NaCl and 2.9% (w/v) n-dodecyl-β-D-maltoside (DDM) to a final detergent:protein weight ratio of 2:1. After centrifugation (20,000 x g, 15 min, 4°C) to remove unsolubilized material, the filtered supernatant was loaded onto a POROS GoPure HQ column (Thermo Fisher Scientific, USA) equilibrated in buffer A (30 mM Tris-HCl, pH 7.8, 2 mM $MgCl_2$, 50 mM NaCl and 0.015% (w/v) DDM) on an Äkta purifier (GE Healthcare). After washing the column with 100mM NaCl in buffer A, ATP synthase dimers were eluted with a linear 100 mM to 300 mM NaCl gradient in buffer A. For the final purification step, fractions containing ATP synthase dimers were concentrated in Vivaspin 500 columns with 100,000 molecular mass cutoff and then loaded onto a Superose6 Increase 3.2/30 size exclusion column (GE Healthcare, USA) equilibrated in buffer B (30 mM Tris-HCl, pH 7.8, 2 mM $MgCl_2$, 40 mM NaCl, 0.05% (w/v) DDM) on an Ettan purifier (GE Healthcare, USA). Fractions of pure and active ATP synthase dimers were collected on ice and used directly for cryo-EM specimen preparation. Purity and activity of the sample was analysed by blue-native polyacrylamide gel electrophoresis (BN-PAGE) and in-gel ATP hydrolysis assay (*38*). For two-dimensional polyacrylamide gel electrophoresis (2D SDS-PAGE) (*39*), subunits of the ATP synthase dimer were first resolved on a 10% acrylamide/8M urea gel before being separated in the second dimension in a 16% acrylamide gel.

##### Electron microscopy and image processing

A solution of 3-4 mg/ml purified complex was applied to C-flat 1/1 or 2/1 holey carbon grids (Science Services GmbH) glow-discharged for 45 s at 0.15 mA. Grids were blotted for 4-5 s at blotforce 20 using a Vitrobot, and vitrified in liquid ethane. Electron micrographs were collected on a Titan Krios G2 with Falcon III detector in counting mode, at 300 kV and 75,000 x magnification for a calibrated pixel size of 1.053 Å. 81-frame movies were automatically recorded with a dose of 0.4 $e^{-}Å^{-2}s^{-1}$, using EPU software. Movies were aligned using MotionCor2 within the Relion3-beta wrapper and CTF parameters were calculated using CTFFind4.1.10. Particles were picked using Gautomatch with templates generated by 2D classification of a previous dataset (*3*), low-pass filtering images to 20 Å. The dataset was cleaned using 3D classification in Relion3. Following a refinement of the full dimer with solvent masking, per-particle CTF parameters and beam tilt were refined, followed by Bayesian polishing and a second round of CTF and beam tilt refinement. The resulting 3D reconstruction gave an overall resolution for the dimer of 2.94 Å; masking of the stationary membrane-bound and lower peripheral stalk portions of the complex improved the resolution for this region to 2.69 Å. Symmetry expansion of the dataset using relion_particle_symmetry_expand, followed by c1 refinement of the upper peripheral stalk gave a resolution of 2.75 Å for this portion of the complex. 3D classification of the pre-aligned symmetry-expanded dataset, with T=20 and no shifts, allowed separation of twelve rotational states, with one class being low resolution and another being a mixture of states; further classification of the latter gave three additional rotatary states for a total of 13 classes. Each was refined first with a mask around the selected monomer, and then with a mask around only the $F_1$ head and rotor portion of the complex. The classes were grouped according to the major rotary state they represent, and these were refined as for the substates. All steps described above were carried out in Relion3. The polished, CTF-refined dataset was imported into cryoSPARC (*40*) and refined using non-uniform refinement with automated masking; this map was used to assist manual building of the less ordered portions of *ASA3* and *ASA9*. For all rotary states and substates, composite maps were generated from the dimer-masked and $F_1$+rotor-masked refined maps using phenix.combine_focused_maps (*41*).

##### Map and model analysis

Models of subunits *a*, *OSCP*, *ε*, and *ASA1-10* were built manually in Coot (*42*). Models of *α, β,γ,δ,* and *c* were refined manually in Coot based on homology models calculated using the ModWeb server (*43*) on templates 5DN6, 5DN6 (*44*), 2V7Q(*45*), 3ZIA(*46*), and 2XND(*47*), respectively. Models were refined by real-space refinement with simulated annealing in Phenix (*48*) and checked manually before deposition. Model quality statistics are as given by the pdb validation server and are summarized in Table S3. For comparison of rotary states and substates, maps were aligned to a map of $F_o$ and the lower peripheral stalk (EMD-xxxx). Models of rotary states and substates were aligned using the Matchmaker tool of USCF Chimera (*49*), fitting to the *a*-subunit. Figures were made using UCSF Chimera and ChimeraX (*50*).

**_Polytomella_ sp. genomic sequence**
Genomic DNA was purified following a published protocol (*51*) with modifications. Briefly, *Polytomella sp.* cells were harvested in their logarithmic phase and resuspended in solubilisation buffer containing 100 mM Tris-HCl, pH 8.0, 40 mM EDTA, 100 mM NaCl, 2% (w/v) Lauryl sarcosine, 1% (w/v) SDS, 20 µg/ml RNase A and 0.9 mg/ml protein kinase K. After 1h at 4°C, DNA was extracted twice with phenol: chloroform: isoamyl alcohol 25:24:1 and the aqueous phase was ethanol-precipitated for 1h on ice. Genomic DNA was further purified by 4M LiCl and 7.5M ammonium acetate before being ethanol-precipitated again and finally washed twice with water. Genomic sequencing was carried out by Microsynth AG with 50-fold sequence coverage. The genome size was ~50Mb. A six-frame translation of the sequence was used as a query database for mass spectrometry results.

**Mass spectrometry**
*LC-MS/MS analyses.* Tryptic digestion of proteins in solution or excised from 2D gels was performed according to published procedures (*52*). Proteolytic digests were loaded by nano-HPLC (Dionex RSLCnano) on reverse-phase columns (trapping column: Acclaim PepMap c18, particle size 2µm, L=20mm; analytical column: Acclaim PepMap c18, particle size 2 µm, L=50 cm; Thermo Fisher Scientific) and eluted in organic phase gradients (Buffer A: 95% $H_2O$, 5% DMSO, 0.1% formic acid; Buffer B: 80% acetonitrile, 15% $H_2O$, 0.1% formic acid). Typically, gradients were ramped from 4% to 48% buffer B in 80 minutes at a 300nl/min flow rate. Peptides eluting from the column were ionised online using a Nanospray Flex Ion source and analysed in an Orbitrap Elite or Q Exactive Plus mass spectrometer. Mass spectra were acquired over the 350-1,600 m/z range at a resolution of 120,000, and sequence information was acquired by computer-controlled, data-dependent automated switching to MS/MS mode in response to collision energies based on mass and charge state of the candidate ions.

*Data processing.* Data sets were processed with the Proteome Discoverer software package (version 2.1.0.81). Spectra were internally recalibrated on autoproteolytic trypsin fragments when applicable. Proteins were identified by matching the derived mass lists against a customized *Polytomella* database (combination of NCBI nr *Polytomella*, a full six-frame translation of a data set acquired by whole-genome shotgun sequencing (Microsynth AG, Balgach, CH) and a list of common contaminants with the program Sequest (Thermo Fisher Scientific, Bremen). In general, a mass tolerance of 10 ppm for parent ion spectra and 0.6 Da for fragment ion spectra, two missed cleavages, oxidation of Met (dynamic modification), acetylation of the protein N-terminus (dynamic modification) and carbamidomethyl-cysteine (fixed modification) were selected as matching parameters in the search program. Results were evaluated using a percolator node (high confidence q-value, FDR < 0.01) to exclude false positives. Proteome data have been uploaded to the PRIDE online repository (*37*).

**Protein sequence analysis**
Homologues of the ASA10 sequence were found by BLAST searches of the NCBI (*53*) and Phytozome (*54*) databases. Sequence alignments were carried out using Clustal Omega (*55*) and formatted using JalView (*56*).

### Figure S1

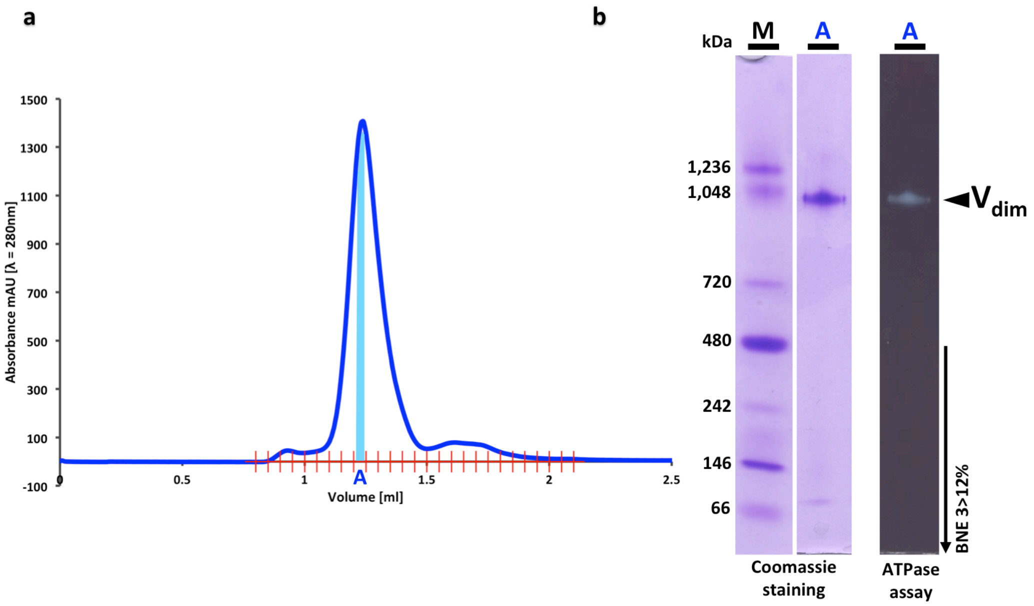

**Figure S1. Protein purification and activity. a)** Purification of mitochondrial ATP synthase dimer from *Polytomella* sp. The light blue bar indicates peak fractions used for cryo-EM grid preparation. **b)** Polyacrylamide gel electrophoresis of the *Polytomella* ATP synthase dimer ($V_{dim}$). M, markers; A, purified gel filtration fractions. An in-gel ATP hydrolysis assay (dark lane on the right) produces a white lead phosphate precipitate, indicating that the dimer is active.

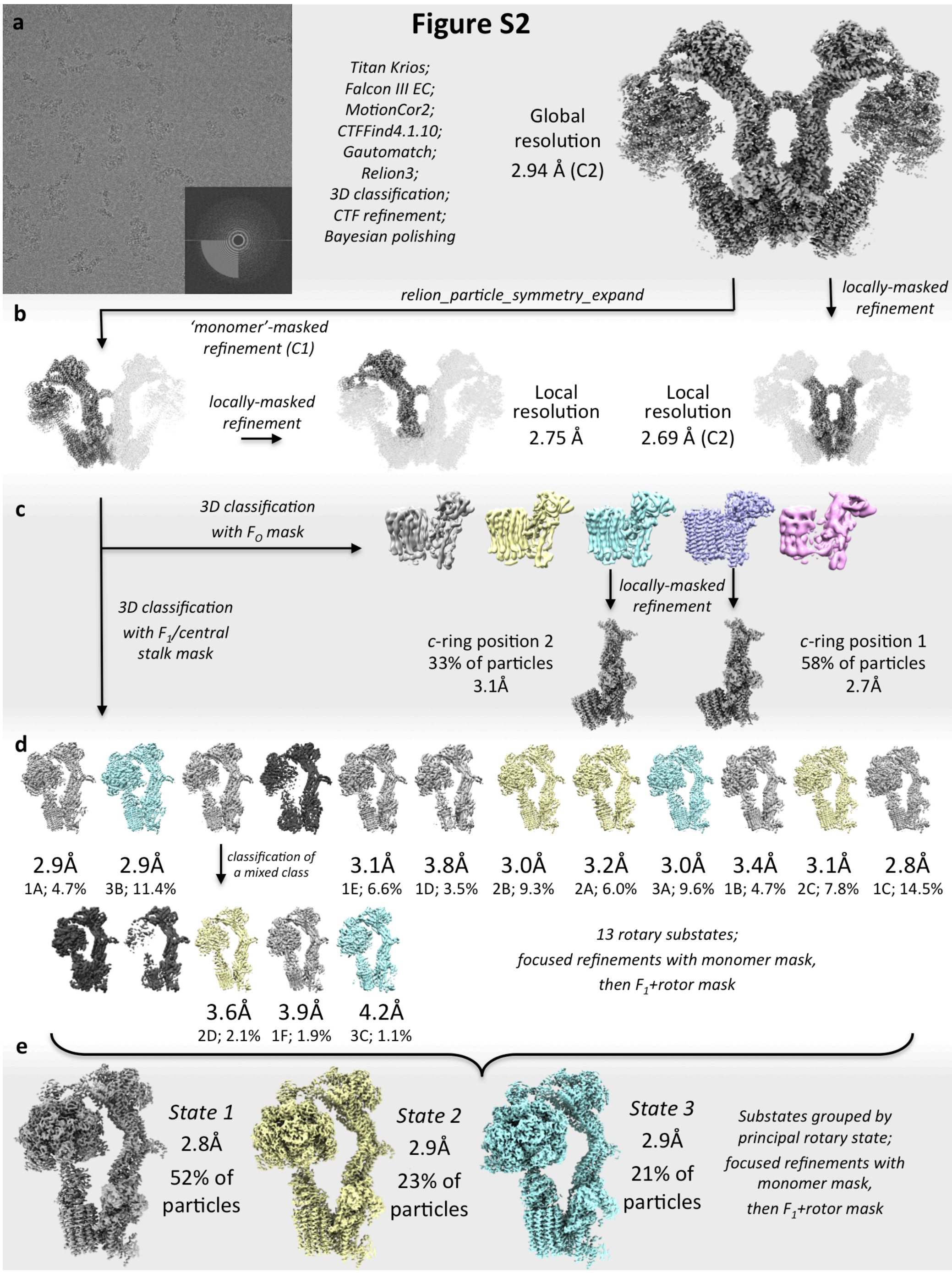

**Figure S2. Cryo-EM image processing scheme. a)** Representative motion-corrected cryo-EM image of *Polytomella* ATP synthase in vitrified buffer, displayed with four-fold Fourier cropping. The Fourier transform of the original image (inset) indicates Thon rings to 3.6 Å resolution. Data were collected with a Krios G2 electron microscope equipped with a Falcon III detector in electron-counting mode at an electron flux of 0.4 $e^{-}Å^{-2}s^{-1}$. Data were collected automatically using EPU software (Thermo Fisher, USA). In total, 18,065 81-frame movies were recorded and aligned with MotionCor2 (*57*). Defocus was estimated with CTFFind4.1.10 (*58*). 735,197 particles were picked with Gautomatch (Dr. Kai Zhang, MRC-LMB) using 20Å low-pass filtering. Further processing was done in Relion3 (*59*). The best 388,670 particles were selected by 3D classification. Bayesian polishing and two rounds of CTF refinement were carried out. 3D refinement resulted in an initial dimer map with 2.94 Å global resolution, with C2 symmetry applied. **b)** Continuing the 3D refinement with a mask applied to the membrane regions and lower peripheral stalk of the dimeric complex gave a resolution of 3.69 Å for this stationary portion of the complex. Symmetry expansion of the particle dataset allowed for independent classification of the rotary state of each monomer in the dimeric complex, and all subsequent steps were carried out on the symmetry-expanded dataset. A focused refinement of the upper portion of the peripheral stalk generated a map with a local resolution of 2.75 Å. **c)** Applying a local mask to the $F_o$ region, the particles were sorted by 3D classification in order to separate distinct positions of the *c*-ring rotor with respect to the stator. Because the $F_1$ head and central stalk were not included in the mask, this classification may blend different $F_1$ rotary states with closely similar *c*-ring positions (given its ten-fold symmetry). Focused refinements of $F_o$ and lower peripheral stalk gave two maps of this region at 2.7 and 3.1 Å resolution. The states differ by a ~13° rotation of the *c*-ring with respect to the membrane-bound stator. **d)** Starting with pre-aligned particles, a mask was applied to $F_1$ and the central stalk to sort rotary substates by 3D classification, using a value of T=20 without particle alignment for the first 25 iterations. Of the twelve classes, one class that contained a mixture of different rotary states was subclassified into five further classes. Excluding three classes of lower resolution (dark gray) presumably containing minor and unresolvable mixed states, we resolved thirteen rotary substates in total, which are colored according to the corresponding primary rotary state they derive from. Each substate was refined with a mask around the monomer, and the refinement was continued with a mask focussing on the $F_1$ head and rotor to obtain a higher resolution map of these regions. The resolution of substates was limited by the number of particles in each class. **e)** Rotary substates were regrouped into three primary rotary states 1, 2 and 3 and refined as for the substates.

**Figure S3**

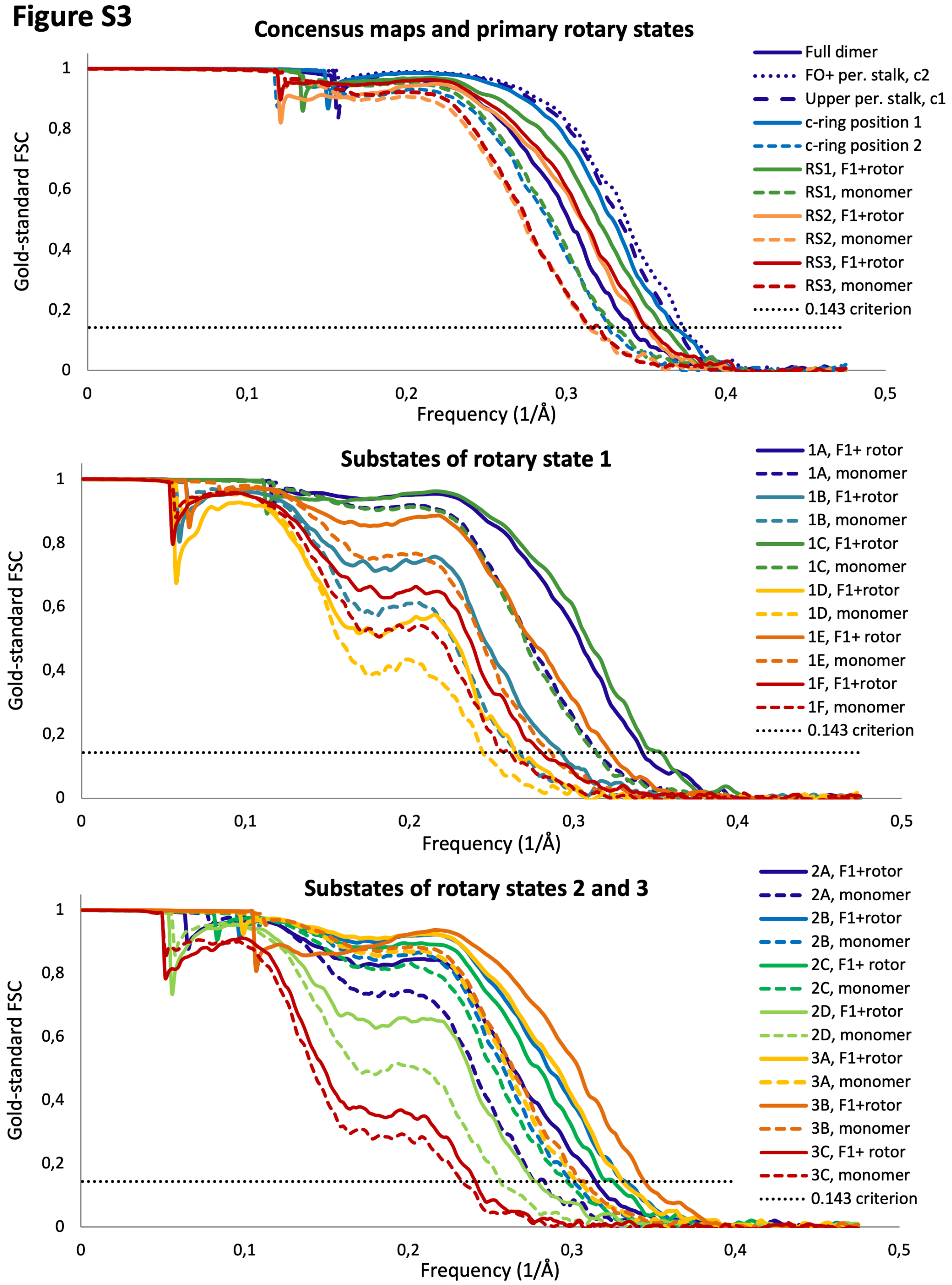

**Figure S3. Corrected FSC curves for post-processing runs of all maps in this work.** **a)** Curves for maps of $F_o$ and peripheral stalk, *c*-ring positions and primary rotary states. **b,c)** Curves for refined rotary substates.

### Figure S4

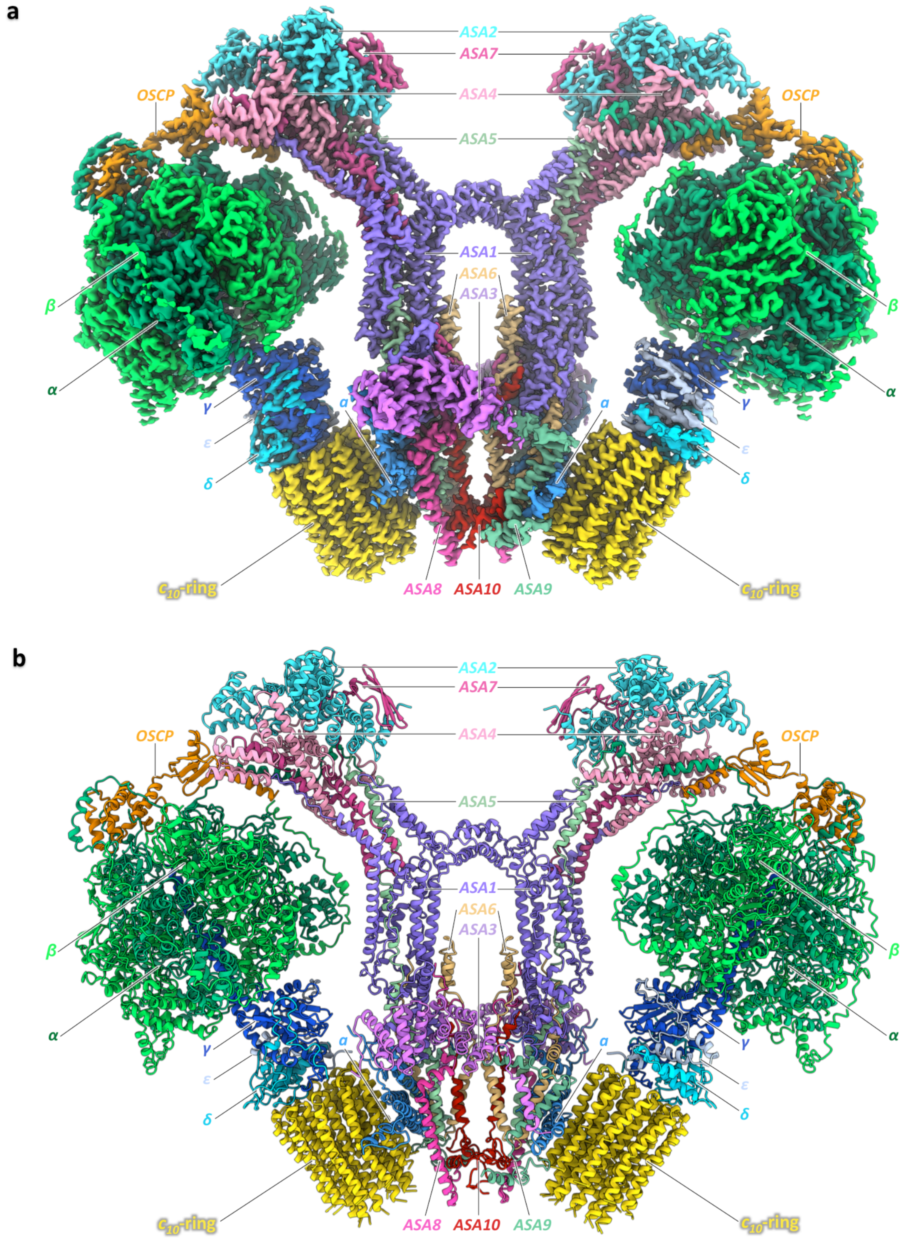

**Figure S4. Atomic model of *Polytomella* ATP synthase dimer. a)** 2.7 to 2.8 Å resolution composite dimer map colored by subunit, as in Figure 1. All subunits are seen from both sides. **b)** Ribbon diagram of atomic model, with subunits colored as in (a) and Figure 1.

#### Figure S5

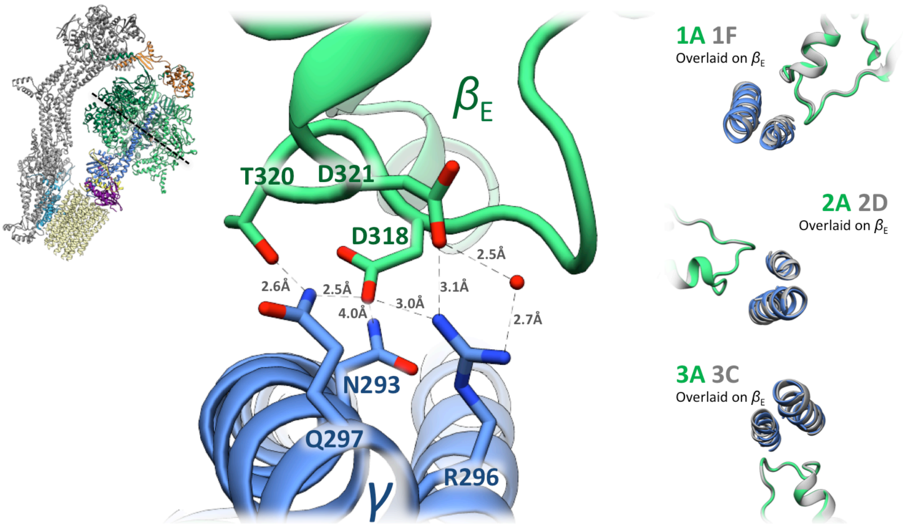

**Figure S5. Interaction of central stalk and $\beta$-subunit catch loop**. Acidic and polar sidechains in the catch loop of subunit $\beta_E$ (green) interact with Asn293 and Gln297 of the $\gamma$ subunit (blue). The interaction with $\alpha$Arg269 includes a resolved water molecule. The inset on the left indicates the section plane. The insets on the right show the first and last substate of each primary rotary state, overlaid with respect to the $\beta_E$ subunit using matchmaker in UCSF Chimera. The interaction of $\beta_E$ and $\gamma$ is essentially unchanged in all rotary substates.

#### Figure S6

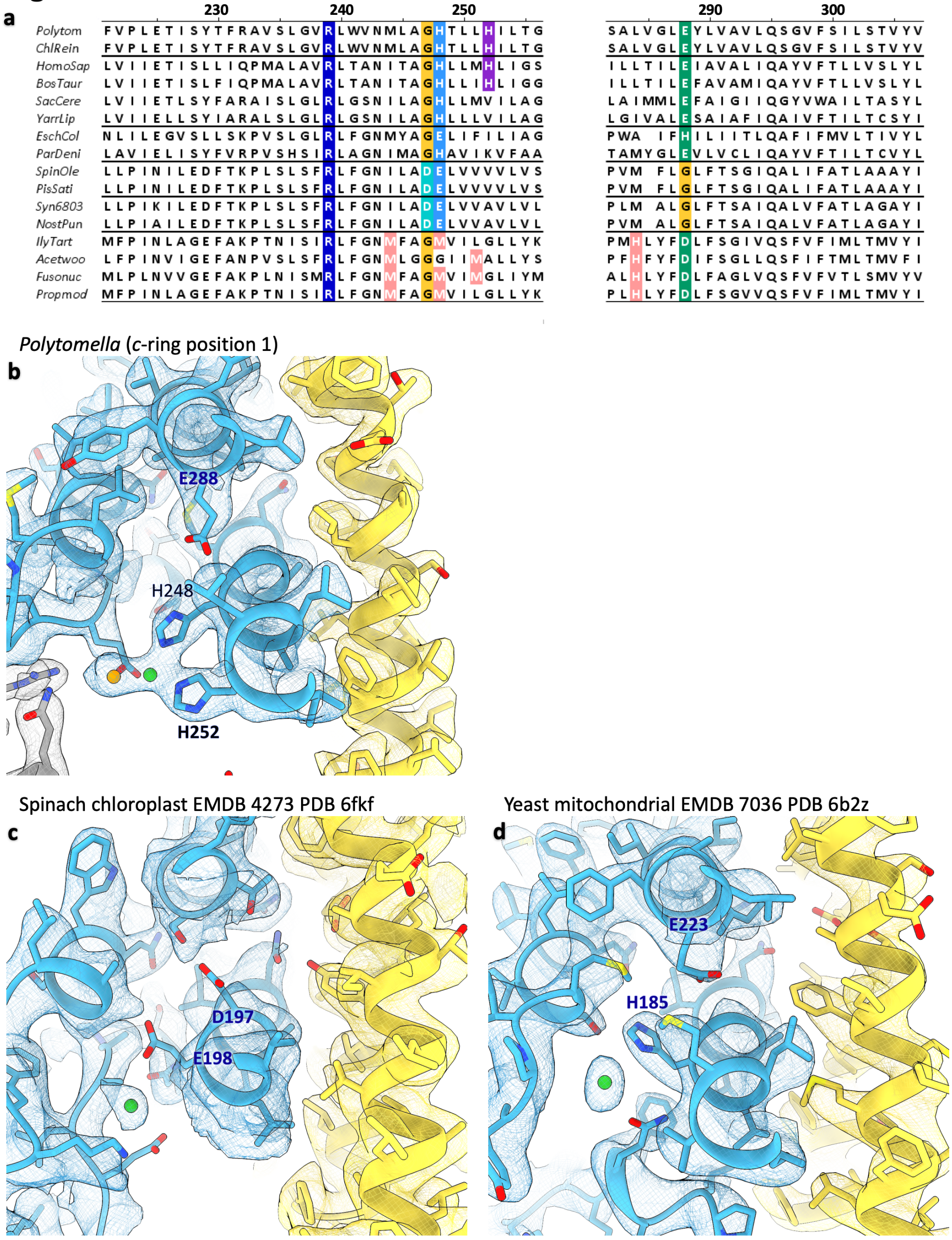

**Figure S6. Conserved ion binding site in the proton access channel of subunit *a*. a)** Sequence alignment of critical regions of subunit *a* helices H5 and H6. The critical *a*Arg239 (navy) separates the aqueous half-channels. In all known mitochondrial and $H^+$-translocating bacterial ATP synthases, positions 248 (light blue) and 288 (dark green) contain one His and one Glu residue; mutation of these residues in *E. coli* abolishes or nearly abolishes both ATP synthesis and $F_o$-mediated proton permeability. Double mutants in which the His and Glu positions are exchanged are more active than either single mutant. A conserved Gly residue (yellow) adjacent to *a*His248 appears to enhance flexibility of *a*H5 in response to *c*-ring pressure. Chloroplast and cyanobacterial ATP synthases have a Glu in the position equivalent to *a*H248 (light blue), while an adjacent Asp residue (cyan; *a*D197 in Spinach chloroplast) appears to be the structural equivalent of *a*E288, being well-positioned to pass protons to the *c*-ring glutamate. The lower four sequences of bacterial sodium-translocating ATPases show clear differences. Potential metal-binding residues located near the His-Glu pair are highlighted in pink. Accession codes are as follows: Chlorophycean algae: *Polytomella* sp. *Pringsheim* 198.80 FN689527.1, *Chlamydomonas reinhardtii* XP_001689492.1; Other mitochondrial ATP synthases: *Homo sapiens* YP_003024031.1, *Bos taurus* YP_209210.1, *Saccharomyces cerevisiae* NP_009313.1, *Yarrowia lipolytica* NP_075433.2; Bacterial ATP synthases: *Escherichia coli* WP_119204347.1, *Paracoccus denitrificans* WP_011749153.1; Chloroplast ATP synthases:*Spinacia oleracea* NP_054920.1, Pisum sativum YP_003587564.1; cyanobacterial ATP synthases: Synechocystis sp. PCC 6803 P27178.1, Nostoc punctiforme PCC

73102 B2J053.1; $Na^+$-translocating ATP synthases: *Ilyobacter tartaricus* AAM94907.1, *Acetobacterium woodii* WP_014354628.1, *Fusobacterium nucleatum* WP_011016339.1, *Propiogenium modestum* P21903.1.
**b)** A strong non-peptide density observed in the proton access channel of *Polytomella* ATP synthase, pictured here for *c*-ring position 1 (see Figure 5 g) is visible in an equivalent position in the lower-resolution cryo-EM maps of spinach chloroplast ATP synthase (*13*) (**c**) and yeast mitochondrial ATP synthase (*35*) (**d**), suggesting an essential, conserved role in proton translocation. In both cases, the metal ion (green sphere) has been added to the model to account for the unassigned density. Maps within a given panel are rendered at the same threshold value.

### Figure S7

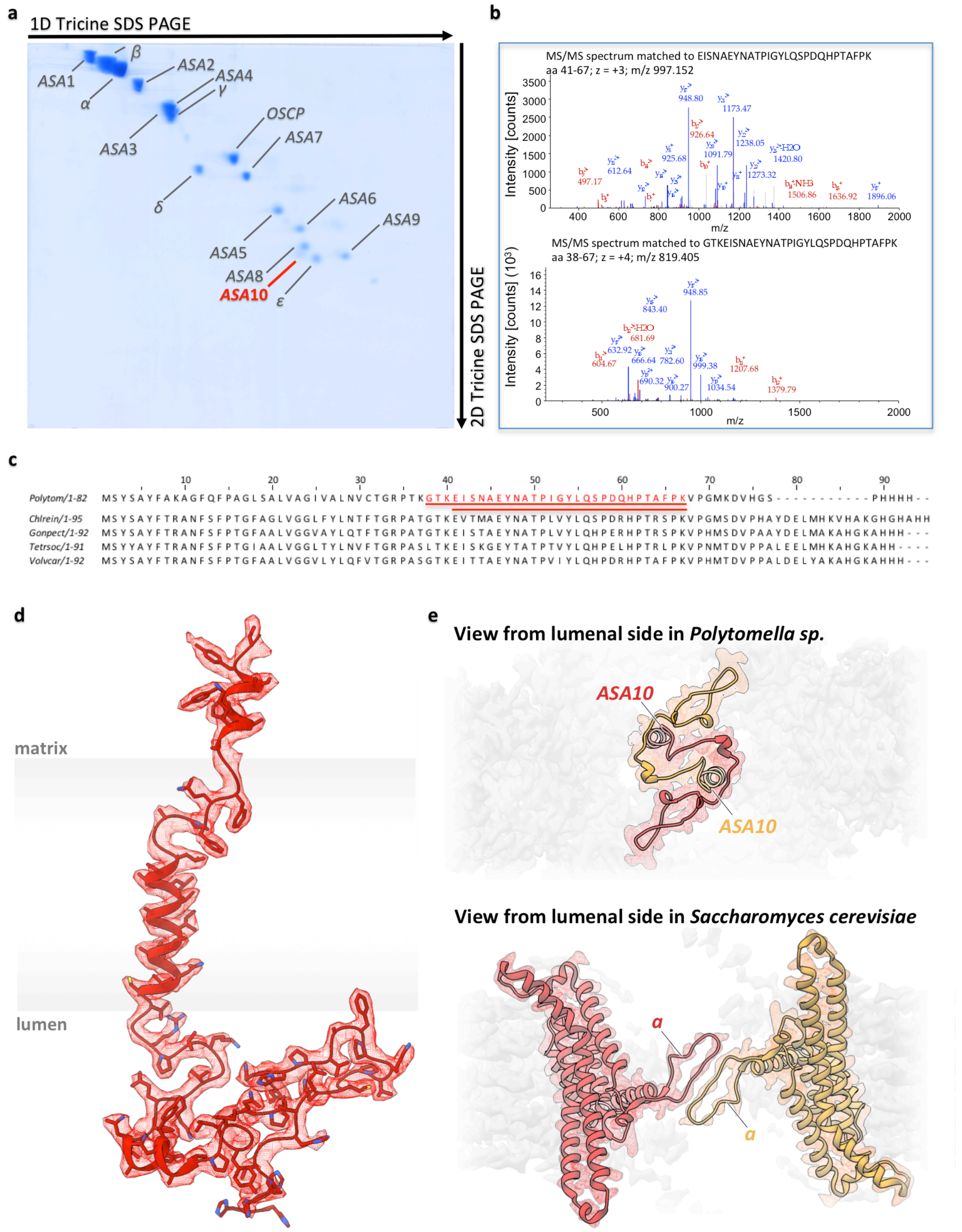

**Figure S7. Subunit *ASA10*. a)** Identification of the new *ASA10* subunit of *Polytomella* ATP synthase by two-dimensional polyacrylamide gel electrophoresis (2D SDS-PAGE) and mass spectrometry. **b)** Representative MS/MS spectra of peptides matched to *ASA10* sequence for indicated amino acid positions. Peaks matched to the peptide sequence are red (b ion series) or blue (y ion series); unassigned peaks are gray. **c)** Complete sequence of *ASA10* with residues identified by mass spectrometry highlighted. BLAST searches and alignment against NCBI (*53*) and Phytozome (*54*) databases revealed homologous unassigned sequences in the proteome of seven chlorophycean algae, of which five are shown: *Chlamydomonas reinhardtii* PNW75022.1, *Gonium pectorale* KXZ41994.1, *Tetrabaena socialis* PNH07028.1, *Volvox carteri* Vocar.0006s0412.1. **d)** Cryo-EM map with fitted atomic model of subunit *ASA10*, indicating its position in the membrane (gray shading). **e)** *ASA10* induces tight interactions between *Polytomella* ATP synthase monomers in the membrane (above). The smaller contact region in the *S. cerevisiae* $F_o$ dimer (*35*) shown below for comparison accounts for the lesser stability of the yeast complex. In the even less stable mammalian ATP synthase dimers, this small protein-protein contact is absent (*1*).

### Figure S8

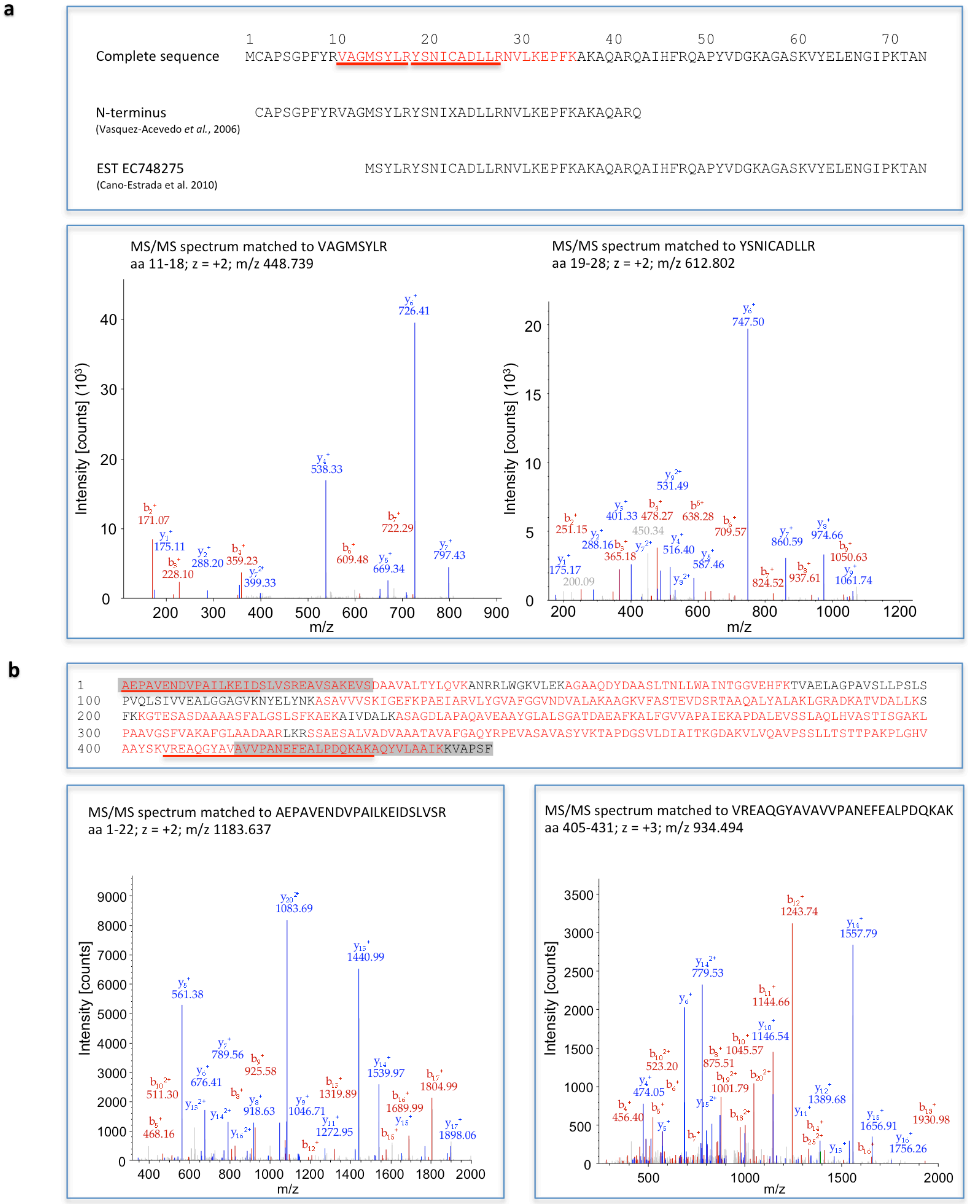

**Figure S8. Biochemistry and mass spectrometry of subunits *ASA2* and $\varepsilon$. a)** Above: Complete sequence of subunit $\varepsilon$ (*60*, *61*) with marked sequence determined by MS. Below: Example MS/MS spectra of selected peptides matched to subunit $\varepsilon$ for amino acid positions indicated. Peaks matched to the peptide sequence are red (b ion series) or blue (y ion series); unassigned peaks are gray. **b)** Above: Complete sequence of subunit *ASA2*. Amino acid segments identified in LC-MS/MS experiments are indicated in red. Red bars indicate selected N- and C-terminal peptides shown in the MS spectra. Newly identified N-and C-terminal segments are shaded in gray. Below: Example MS/MS spectra of selected peptides matched to *ASA2* for indicated amino acid positions. Peaks successfully matched to the peptide sequence indicated in red (b ion series) and blue (y ion series); unassigned peaks indicated in gray.

**Figure S9**

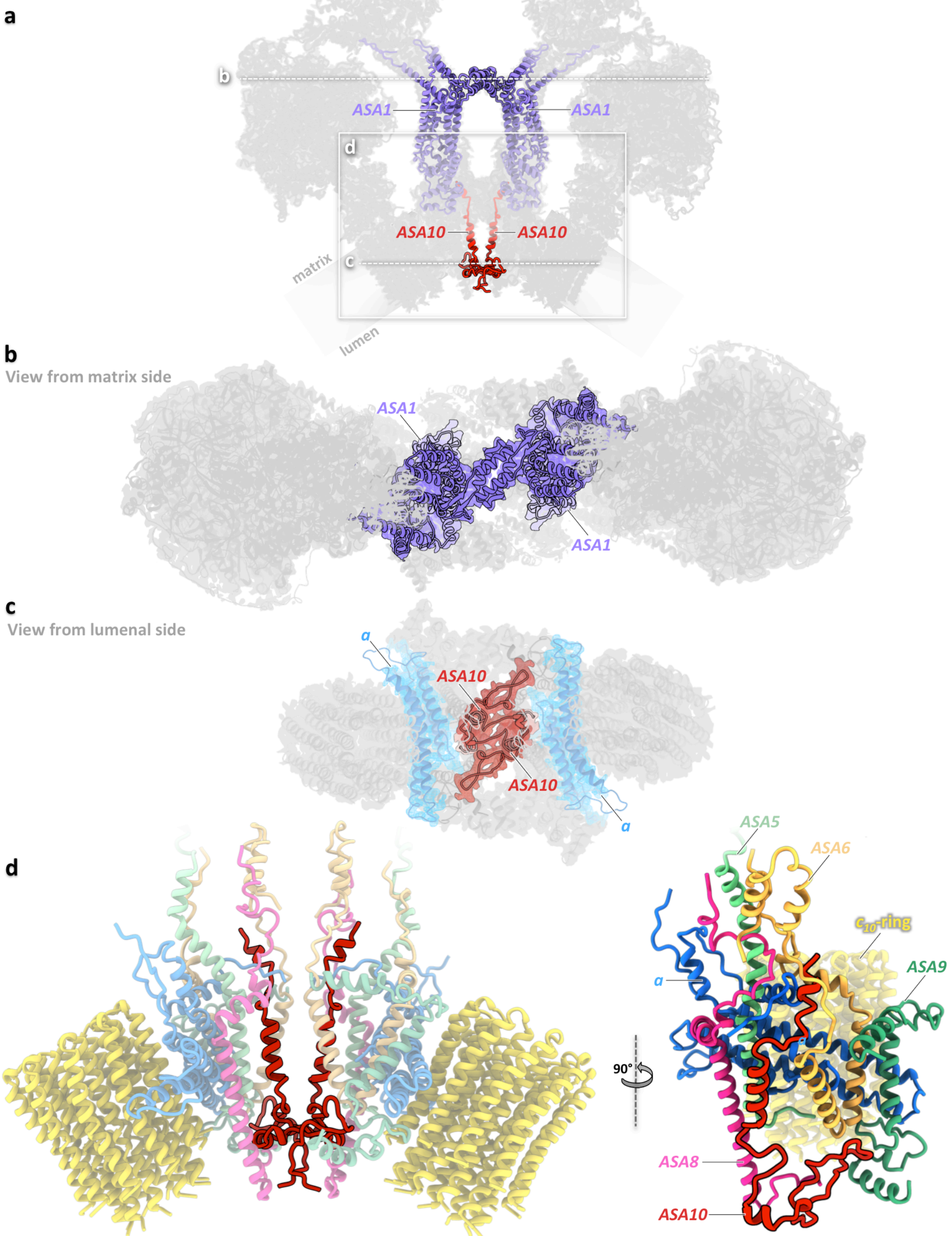

**Figure S9. Protein interactions at the *Polytomella* ATP synthase dimer interface. a)** Positions of subunits *ASA1* (purple) and *ASA10* (red) in the *Polytomella* dimer (light grey). **b)** Two pairs of short $\alpha$-helices of *ASA1* form a four-helix bundle as a protein bridge between the peripheral stalks about 75 Å above the membrane surface on the matrix side. **c,d)** Subunits *ASA10* of the two monomers are closely intertwined, establishing tight contacts between the two $F_o$ regions. The *Polytomella* $F_o$ complex consists of 7 subunits: *ASA5, ASA6, ASA8, ASA9, ASA10,* subunit *a* and the $c_{10}$-ring rotor.

**Movie S1. Three-dimensional map of Polytomella ATP synthase dimer.** One monomer is gray, the other monomer map is colored by subunit, as in Figure 1.

**Movie S2. Morph of 13 rotary substates of the *Polytomella* ATP synthase monomer**. The *c*-ring (yellow) and central stalk (subunits $\gamma$; blue; $\delta$, cyan and $\varepsilon$, pale blue) together rotate in three roughly equal ~120° steps. The $F_1$ head moves with them for the first ~30° of each step. $\alpha$, dark green; $\beta$, bright green. The two-domain *OSCP* subunit (orange) works as a hinge between the moving $F_1$ head and the stationary peripheral stalk (grey).

**Movie S3. Section through morph of Movie S2 at the level of nucleotide binding sites in the $F_1$ head**. Catalytic sites of subunits $\beta$ (bright green) are orange. Binding sites for the structural ATP in the three $\alpha$-subunits (dark green) are red. The $\beta$ catch loop is drawn in purple. Central stalk subunit $\gamma$; blue; peripheral stalk, gray.

**Movie S4. Hinge movement of the two-domain *OSCP* subunit (orange).** The proximal $\alpha$-helical *OSCP* domain on the right is attached to the $F_1$ head by the N-terminal extensions of two of the three $\alpha$ -subunits (dark green). The distal $\beta$-sheet *OSCP* domain is attached to the peripheral stalk (gray) by interaction of one *OSCP* helix and the N-terminal extension of two of the three $\alpha$ -subunits.

**Movie S5. Extensions of subunits $\alpha$ and $\beta$ in *Polytomella* ATP synthase**
The polypeptide sequence of subunits $\alpha$ and $\beta$ in mitochondrial $F_1F_o$ ATP synthases of chlorophycean algae (including *Polytomella* sp.) have characteristic extensions at their N and C termini. The ~15 residue N-terminal extension of subunit $\alpha$ interacts with *OSCP* (see Figure 4 and Movie S4). The ~60-residue C-terminal extension of the $\beta$-subunit wraps around the outside of the neighboring subunit $\alpha$ in the $F_1$ head. Both extensions would render the chlorophycean $F_1F_o$ complex more stable than that of mammalian or fungal mitochondrial ATP synthases.

**Table S1. EM Statistics**

| | |
|---|---|
| **Data collection** | |
| Electron Microscope | Titan Krios G2 |
| Camera | Falcon III (electron-counting mode) |
| Data collection software | EPU (Thermo Fisher) |
| Voltage | 300 kV |
| Nominal magnification | 75,000 x |
| Calibrated physical pixel size | 1.053 Å |
| Total exposure | 35 $e^{-}Å^{-2}$ |
| Exposure rate | 0.4 $e^{-}Å^{-2}s^{-1}$ |
| Number of frames | 81 |
| Defocus range | −0.4 to −5 µm (95% between -0.8 and -2.2 µm) |
| **Image Processing** | |
| Motion correction software | MotionCor2 |
| CTF estimation software | CTFFind4.1.10 |
| Particle selection software | Gautomatch |
| Micrographs used | 18,065 |
| Particles selected | 735,197 |
| Cassification and refinement software | Relion3 |
| Particles contributing to final dataset | 388,670 |
| **Model Building** | |
| Modeling software | ModWeb, Coot |
| Refinement software | Phenix (phenix.real_space_refine) |

**Table S2. Map identifiers and statistics**

| Description | EMDB ID | Associated model | Resolution (0.143 criterion) | Applied b-factor | Number of particles | Applied symmetry |
|---|---|---|---|---|---|---|
| Full dimer | | | 2.94 Å | -69 | 388,670 | C2 |
| $F_o$/peripheral stalk | | | 2.69 Å | -61 | 388,670 | C2 |
| Upper peripheral stalk | | | 2.75 Å | -66 | 777,340 | C1 |
| c-ring position 1 | | | 2.73 Å | -57 | 536,992 | C1 |
| c-ring position 2 | | | 3.08 Å | -51 | 136,422 | C1 |
| Primary Rot. State 1 | | | Composite map of EMDx and EMDx | | | |
| RS1 monomer | | | 3.04 Å | -65 | 400,918 | C1 |
| RS1 $F_1$+rotor | | | 2.78 Å | -56 | 400,918 | C1 |
| Primary Rot. State 2 | | | Composite map of EMDx and EMDx | | | |
| RS2 monomer | | | 3.20 Å | -68 | 179,651 | C1 |
| RS2 $F_1$+rotor | | | 2.86 Å | -54 | 179,651 | C1 |
| Primary Rot. State 3 | | | Composite map of EMDx and EMDx | | | |
| RS3 monomer | | | 3.18 Å | -64 | 163,259 | C1 |
| RS3 $F_1$+rotor | | | 2.87 Å | -45 | 163,259 | C1 |
| Rot. Substate 1A | | | Composite map of EMDx and EMDx | | | |
| RS1A monomer | | | 3.20 Å | -62 | 124,537 | C1 |
| RS1A $F_1$+rotor | | | 2.94 Å | -44 | 124,537 | C1 |
| Rot. Substate 1B | | | Composite map of EMDx and EMDx | | | |
| RS1B monomer | | | 3.74 Å | -67 | 72,402 | C1 |
| RS1B $F_1$+rotor | | | 3.44 Å | -46 | 72,402 | C1 |
| Rot. Substate 1C | | | Composite map of EMDx and EMDx | | | |
| RS1C monomer | | | 3.20 Å | -59 | 112,810 | C1 |
| RS1C $F_1$+rotor | | | 2.84 Å | -45 | 112,810 | C1 |
| Rot. Substate 1D | | | Composite map of EMDx and EMDx | | | |
| RS1D monomer | | | 4.11 Å | -71 | 27,039 | C1 |
| RS1D $F_1$+rotor | | | 3.80 Å | -52 | 27,039 | C1 |

| Description | EMDB ID | Associated model | Resolution (0.143 criterion) | Applied b-factor | Number of particles | Applied symmetry |
|---|---|---|---|---|---|---|
| Rot. Substate 1E | | | Composite map of EMDx and EMDx | | | |
| RS1E monomer | | | 3.51 | -65 | 51,482 | C1 |
| RS1E $F_1$+rotor | | | 3.12 | -49 | 51,482 | C1 |
| Rot. Substate 1F | | | Composite map of EMDx and EMDx | | | |
| RS1F monomer | | | 3.85 | -59 | 15,099 | C1 |
| RS1F $F_1$+rotor | | | 3.58 | -53 | 15,099 | C1 |
| Rot. Substate 2A | | | Composite map of EMDx and EMDx | | | |
| RS2A monomer | | | 3.64 | -67 | 46,820 | C1 |
| RS2A $F_1$+rotor | | | 3.20 | -49 | 46,820 | C1 |
| Rot. Substate 2B | | | Composite map of EMDx and EMDx | | | |
| RS2B monomer | | | 3.33 | -54 | 72,402 | C1 |
| RS2B $F_1$+rotor | | | 3.01 | -44 | 72,402 | C1 |
| Rot. Substate 2C | | | Composite map of EMDx and EMDx | | | |
| RS2C monomer | | | 3.37 | -62 | 60,429 | C1 |
| RS2C $F_1$+rotor | | | 3.08 | -44 | 60,429 | C1 |
| Rot. Substate 2D | | | Composite map of EMDx and EMDx | | | |
| RS2D monomer | | | 3.92 | -64 | 16,363 | C1 |
| RS2D $F_1$+rotor | | | 3.61 | -45 | 16,363 | C1 |
| Rot. Substate 3A | | | Composite map of EMDx and EMDx | | | |
| RS3A monomer | | | 3.30 | -66 | 74,956 | C1 |
| RS3A $F_1$+rotor | | | 3.03 | -44 | 74,956 | C1 |
| Rot. Substate 3B | | | Composite map of EMDx and EMDx | | | |
| RS3B monomer | | | 3.26 | -58 | 88,303 | C1 |
| RS3B $F_1$+rotor | | | 2.92 | -42 | 88,303 | C1 |
| Rot. Substate 3C | | | Composite map of EMDx and EMDx | | | |
| RS3C monomer | | | 4.32 | -82 | 8,173 | C1 |
| RS3C $F_1$+rotor | | | 4.18 | -67 | 8,173 | C1 |

**Table S3. Model identifiers and quality statistics**

| Description | PDB ID | Associated map | Residues built | RMS bonds Length (Å) | RMS bonds Angles (°) | Ramachandran outliers (%) | Ramachandran favoured (%) | Rotamer outliers (%) | Clashscore | EMRinger score |
|---|---|---|---|---|---|---|---|---|---|---|
| Full dimer | | | | | | | | | | |
| $F_o$/peripheral stalk | | | 1,586 | 0.014 | 1.050 | 0.19 | 94.27 | 0.24 | 3.64 | 4.07 |
| Upper peripheral stalk | | | 1,027 | 0.006 | 0.807 | 0.20 | 95.38 | 0.13 | 5.37 | 3.68 |
| c-ring position 1 | | | 2,325 | 0.010 | 0.942 | 0.22 | 94.89 | 0.17 | 6.04 | 3.15 |
| c-ring position 2 | | | 2,325 | 0.006 | 0.896 | 0.22 | 94.80 | 0.34 | 6.30 | 3.03 |
| Primary Rot. State 1 | | | 7,119 | 0.014 | 1.106 | 0.23 | 94.95 | 0.69 | 6.00 | 3.64 |
| RS1 monomer | | | 2,613 | 0.013 | 1.033 | 0.23 | 94.47 | 0.54 | 3.73 | 3.64 |
| RS1 $F_1$+rotor | | | 4,506 | 0.008 | 0.951 | 0.29 | 95.27 | 0.34 | 5.96 | 3.38 |
| Primary Rot. State 2 | | | 7,119 | 0.013 | 1.058 | 0.18 | 94.50 | 0.41 | 5.54 | 3.50 |
| RS3 monomer | | | 2,613 | 0.012 | 1.023 | 0.27 | 94.55 | 0.20 | 4.49 | 3.42 |
| RS2 $F_1$+rotor | | | 4,506 | 0.010 | 0.996 | 0.13 | 95.18 | 0.36 | 6.15 | 3.51 |
| Primary Rot. State 3 | | | 7,120 | 0.011 | 0.970 | 0.23 | 94.98 | 0.48 | 5.49 | 3.59 |
| RS3 monomer | | | 2,613 | 0.008 | 0.889 | 0.19 | 94.74 | 0.20 | 3.83 | 3.56 |
| RS3 $F_1$+rotor | | | 4,507 | 0.005 | 0.814 | 0.16 | 96.26 | 0.20 | 5.66 | 3.56 |
